# Distinct TAF15 amyloid filament folds define multiple subtypes of FTLD-TAF15

**DOI:** 10.64898/2026.01.12.698957

**Authors:** Stephan Tetter, Nikhil R. Varghese, Alexey G. Murzin, Wouter De Coster, Marleen Van den Broeck, Sigrun Roeber, Jeffrey T. Joseph, Kathy Newell, Rudolf Castellani, Sumit Das, Lee-Cyn Ang, Matthis Synofzik, Jochen Herms, Rosa Rademakers, Bernardino Ghetti, Tammaryn Lashley, Ian R. A. Mackenzie, Manuela Neumann, Benjamin Ryskeldi-Falcon

## Abstract

Neurodegenerative diseases are characterised by the assembly of a limited number of disease-specific proteins into amyloid filaments, which form intracellular inclusions or extracellular deposits in the central nervous system (CNS)^1,2^. We previously found that amyloid filaments of TATA-binding protein-associated factor 15 (TAF15) characterise a subtype of frontotemporal lobar degeneration with FET protein-immunoreactive inclusions (FTLD-FET)^3^, termed atypical FTLD with ubiquitin-positive inclusions (aFTLD-U)^4^, which causes early-onset, rapidly progressive behavioural variant frontotemporal dementia (FTD). However, it was not clear if TAF15 proteinopathy was more widespread in neurodegenerative diseases. Two additional FTLD-FET subtypes have been proposed, neuronal intermediate filament inclusion body disease (NIFID) and basophilic inclusion body disease (BIBD)^5,6^, which have more heterogenous clinical presentations including FTD, motor neuron diseases (MND) and movement disorders. Here, we used electron cryo-microscopy (cryo-EM) to determine a total of 32 amyloid filament structures from the brains of 17 individuals encompassing all three proposed subtypes of FTLD-FET and their diverse clinical presentations. All cases were characterised by TAF15 filaments, in the absence of filaments of the other FET proteins, fused in sarcoma (FUS) and Ewing’s sarcoma (EWS). All three aFTLD-U cases had the previously-reported TAF15 fold^3^. Unexpectedly, we found four distinct TAF15 folds among 11 NIFID cases. Eight of these cases shared a common fold, while the remaining three were each distinct. Furthermore, we found distinct TAF15 folds for each of the three BIBD cases. Neuropathological reassessment of the neocortical TAF15 inclusion pathology of these cases distinguished the NIFID cases with the common fold from the others. Thus, TAF15 filament structures form the basis of a new, expanded classification of FTLD-FET subtypes. Moreover, we discovered a TAF15 Y38C variant in the filament fold of one of the individuals with BIBD. The structure is unable to incorporate wild-type TAF15, despite the individual being heterozygous, suggesting that this variant drives TAF15 filament assembly. This study provides structural and genetic evidence that TAF15 amyloid filaments underlie the diverse group of neurodegenerative diseases currently termed FTLD-FET, which we therefore rename FTLD-TAF15.

## INTRODUCTION

Neurodegenerative diseases are devastating conditions characterised by the progressive loss of neurons. Incurable to date, an improved molecular understanding is paramount to improving diagnosis and treatment. Neuronal loss is preceded by the accumulation of amyloid filaments within intracellular inclusions and, in certain diseases, extracellular deposits in the CNS^1,2^. Amyloid filaments in neurodegenerative diseases are formed by approximately a dozen known proteins, each of which is associated with a specific subset of diseases^2^. Pathogenic variants in the genes encoding these proteins enhance their amyloid assembly, thereby establishing a causal role for this process in disease^1^.

Amyloid filaments are characterised by the stacking of an essentially planar protein fold^7^. This ordered filament core is stabilised by extensive intermolecular hydrogen-bonding networks, including β-sheets and amino acid side chain interactions. Similar to globular protein folds^8^, amyloid core folds can exhibit local structural variation, resulting in filaments containing a mixture of several variants of a single fold^9,10^. The filament core is typically formed by only a portion of the protein and is surrounded by a less ordered ‘fuzzy coat’ of the flanking protein sequences^11^.

Cryo-EM structures of amyloid filament cores from patients’ brains have established that specific filament folds of the proteins tau, α-synuclein and TDP-43 characterise different diseases^2^. This occurs despite the fact that many, likely all^12^, proteins can form a multitude of different amyloid filament folds *in vitro*^7^. These findings suggest that distinct molecular mechanisms underlie different neurodegenerative diseases with amyloid filaments at their core. The filament fold structures also provide an opportunity to design specific tools for the diagnosis and, potentially, treatment of neurodegenerative diseases.

The FET protein family, FUS, EWS and TAF15, are ubiquitous nucleocytoplasmic-shuttling RNA-binding proteins^13^. They comprise an N-terminal low complexity domain (LCD) enriched in glycine, tyrosine, glutamine and serine residues; two RGG motif-rich segments; an RNA recognition motif; a zinc finger domain; and a C-terminal nuclear localisation signal. In health, the FET proteins help to regulate multiple stages of gene expression, including in transcription and RNA maturation.

FTLD-FET is characterised by abundant FET protein-immunoreactive neuronal inclusions^14^. It accounts for approximately 10-20% of FTLD cases, with the remaining cases characterised by TDP-43 (FTLD-TDP) or tau (FTLD-tau) filament inclusions^15^. FTLD-FET was previously referred to as FTLD-FUS, since FUS was the first FET protein for which inclusion immunoreactivity was established^4^, and pathogenic mutations in *FUS* had previously been linked to a rare form of amyotrophic lateral sclerosis (ALS)^16,17^. However, to date, no genetic variation has been associated with FTLD-FET.

Three subtypes of FTLD-FET have been proposed: aFTLD-U, NIFID and BIBD^4–6,18^. NIFID is distinguished by the presence of inclusions composed of all class IV neuronal intermediate filaments^19,20^, which are found in a subset of neurons containing FET protein-immunoreactive inclusions^5,18^. BIBD is distinguished by the presence of large basophilic inclusions^21,22^, which correspond to a subset of FET protein neuronal cytoplasmic inclusions (NCIs)^6,18^. Whereas FET protein-immunoreactive NCIs in aFTLD-U have a distinct compact round morphology, they are more heterogeneous in NIFID and BIBD, including compact round, annular and tangle-like, in addition to dystrophic neurites and neuropil dots^18,23^. In addition, vermiform neuronal intranuclear inclusions are present in aFTLD-U and NIFID but are not observed in BIBD. Clinically, aFTLD-U usually manifests as early-onset, rapidly progressive behavioural variant FTD (bvFTD). In contrast, NIFID and BIBD are more heterogeneous, presenting as FTD, MND and/or extrapyramidal dysfunction^18,23^.

We recently found that TAF15 amyloid filaments with a single fold characterised the FTLD-FET subtype aFTLD-U in four individuals examined^3^. This discovery added TAF15 to the limited number of proteins that form amyloid filaments in neurodegenerative diseases^2^. Against expectations, we did not find any evidence for amyloid filament formation of FUS, or of the third FET protein EWS. The presence, identities and structures of amyloid filaments in the other proposed subtypes of FTLD-FET – NIFID and BIBD – were unknown. In addition, genetic evidence linking TAF15 amyloid filaments with the pathogenesis of FTLD was lacking. Here, we investigated the identities and structures of amyloid filaments in all three proposed subtypes of FTLD-FET, spanning their diverse clinical presentations.

## RESULTS

### TAF15 amyloid filaments characterise aFTLD-U, NIFID and BIBD

We extracted detergent-insoluble proteins from postmortem prefrontal cortex of an additional three individuals with aFTLD-U, 11 individuals with NIFID and three individuals with BIBD (Supplementary Figure 1a and Supplementary Tables 1 and 2), using differential centrifugation in the presence of the detergent *N*-lauroyl-sarcosine (sarkosyl) as in our previous study of aFTLD-U^3^. The individuals encompassed the diverse clinical presentations of FTLD-FET, including FTD, MND and extrapyramidal dysfunction (Supplementary Table 1). Immunoblots confirmed the presence of all three FET proteins in the extracts (Supplementary Figure 1b), consistent with their inherent detergent insolubility^14^.

Cryo-EM imaging revealed abundant amyloid filaments in the extracts from all 17 individuals. We used helical reconstruction in RELION to determine a total of 32 structures of filaments’ ordered cores from the cryo-EM images^24^, with resolutions of between 1.6 Å and 3.7 Å (Supplementary Figure 2 and Supplementary Table 3). These high-resolution structures enabled atomic modelling, thereby unambiguously identifying the protein that formed each filament structure (Figures 1-3 and Supplementary Figures 3-8). The filaments from each case of aFTLD-U, NIFID and BIBD were composed of TAF15, rather than the more widely studied FET protein FUS or the third family member EWS. The ordered cores of all of the filaments were formed by the approximately 100 N-terminal residues of TAF15, comprising the N-terminal half of its LCD. These results strongly suggest that TAF15 amyloid filaments characterise all proposed subtypes of FTLD-FET.

**Figure 1:**
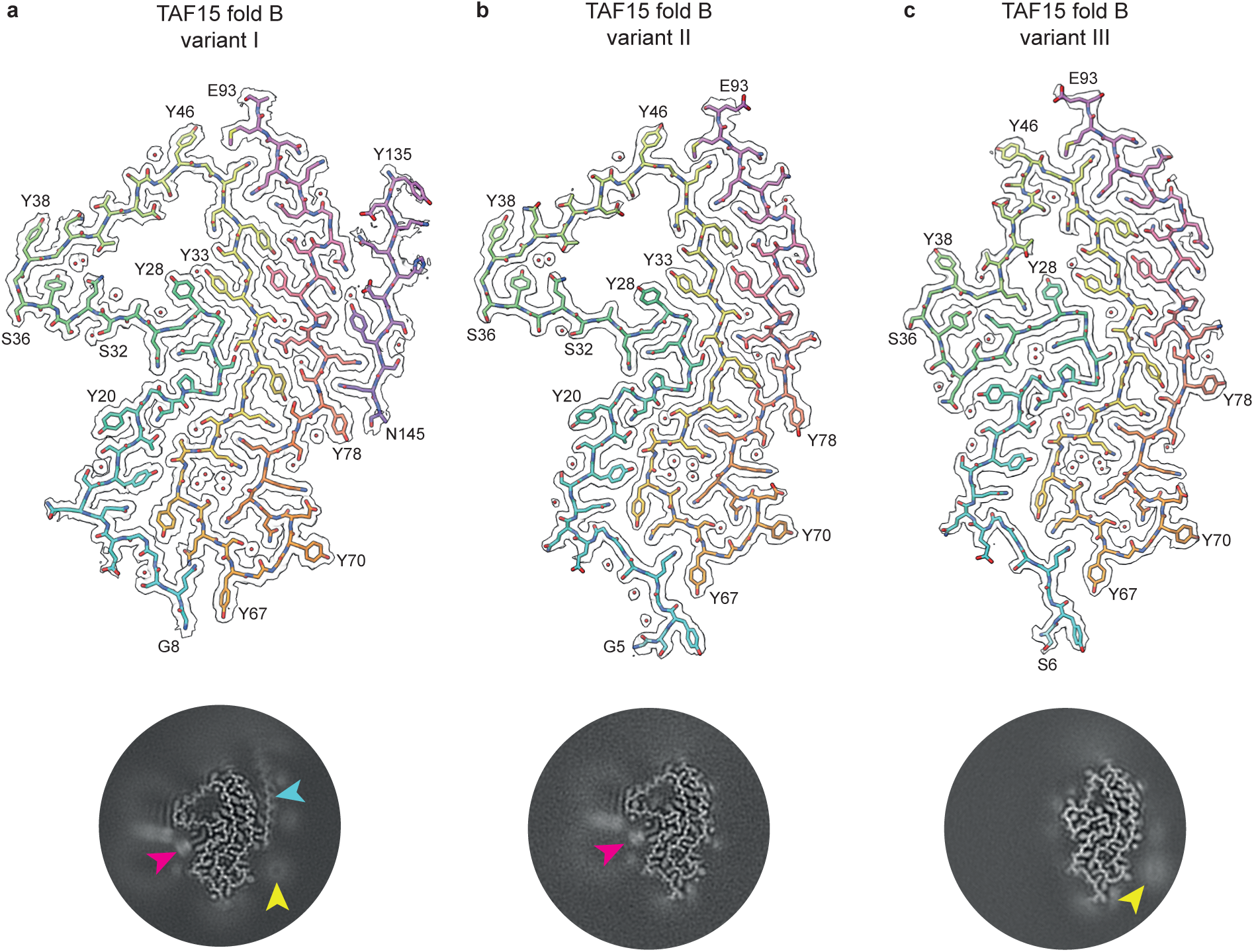
TAF15 filament fold B in NIFID. a-c, Cryo-EM reconstructions (black outlines) and atomic models of variants I **(a)**, II **(b)** and III **(c)** of TAF15 filament fold B, shown for a single TAF15 molecule perpendicular to the helical axis. The cryo-EM reconstructions are shown using a surface zone around the atomic models. The atomic models are colour-coded as a gradient from N– to C-terminal residues. Central slices of the cryo-EM reconstructions centred on the helical axis are shown as inserts. A disconnected TAF15 Y135-N145 peptide (cyan arrow), lipid-like densities (magenta arrow) and large round disconnected densities (yellow arrow) are highlighted.

**Figure 2:**
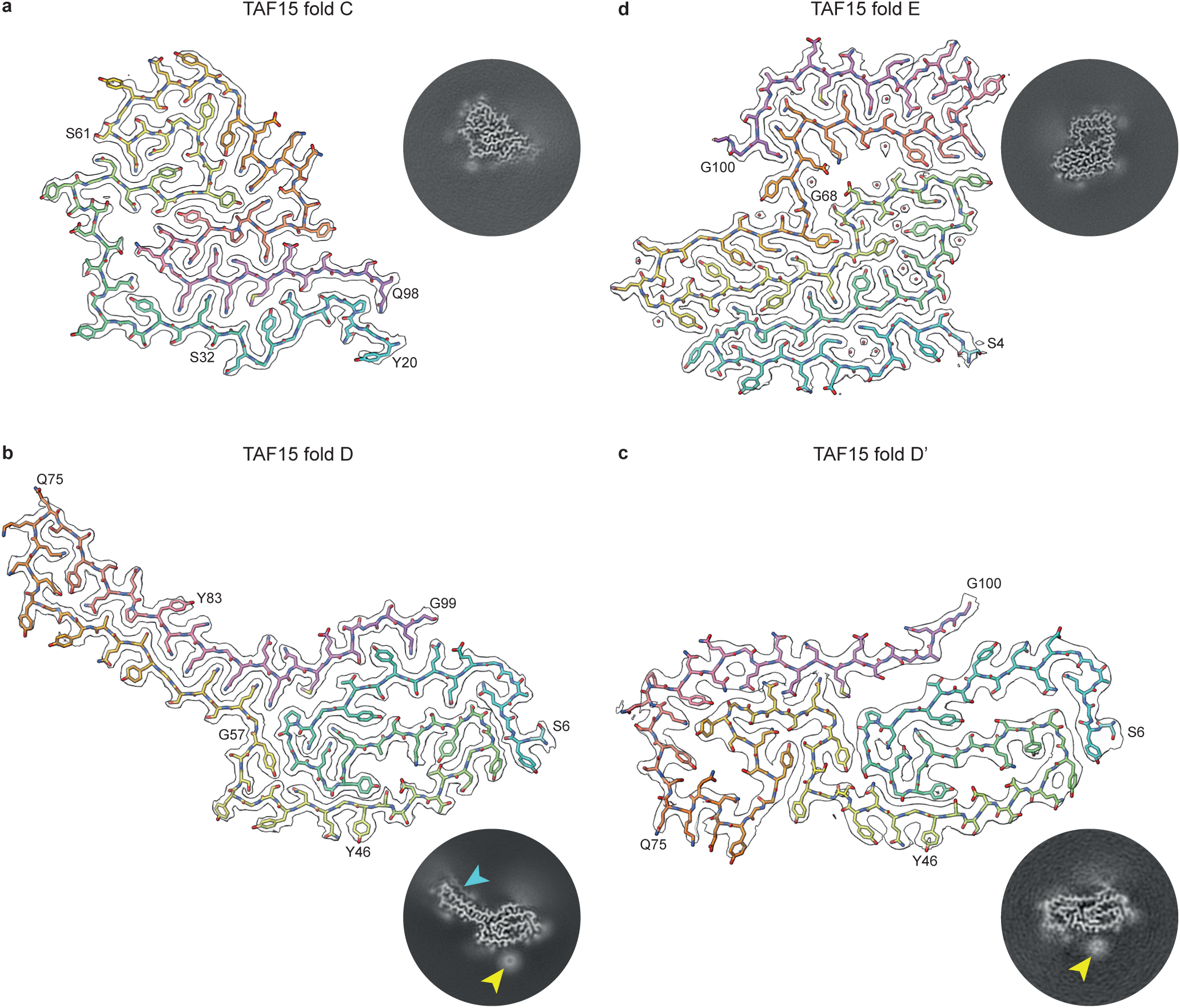
TAF15 filament folds C, D and E in NIFID. a-d, Cryo-EM reconstructions (black outlines) and atomic models of TAF15 filament fold C **(a)**, fold D **(b)**, fold D’ **(c)** and fold E **(d)**, shown for a single TAF15 molecule perpendicular to the helical axis. The cryo-EM reconstructions are shown using a surface zone around the atomic models. The atomic models are colour-coded as a gradient from N– to C-terminal residues. Central slices of the cryo-EM reconstructions centred on the helical axis are shown as inserts. A disconnected peptide (cyan arrow) and large round disconnected density (yellow arrow) associated with folds D and D’ are highlighted.

**Figure 3:**
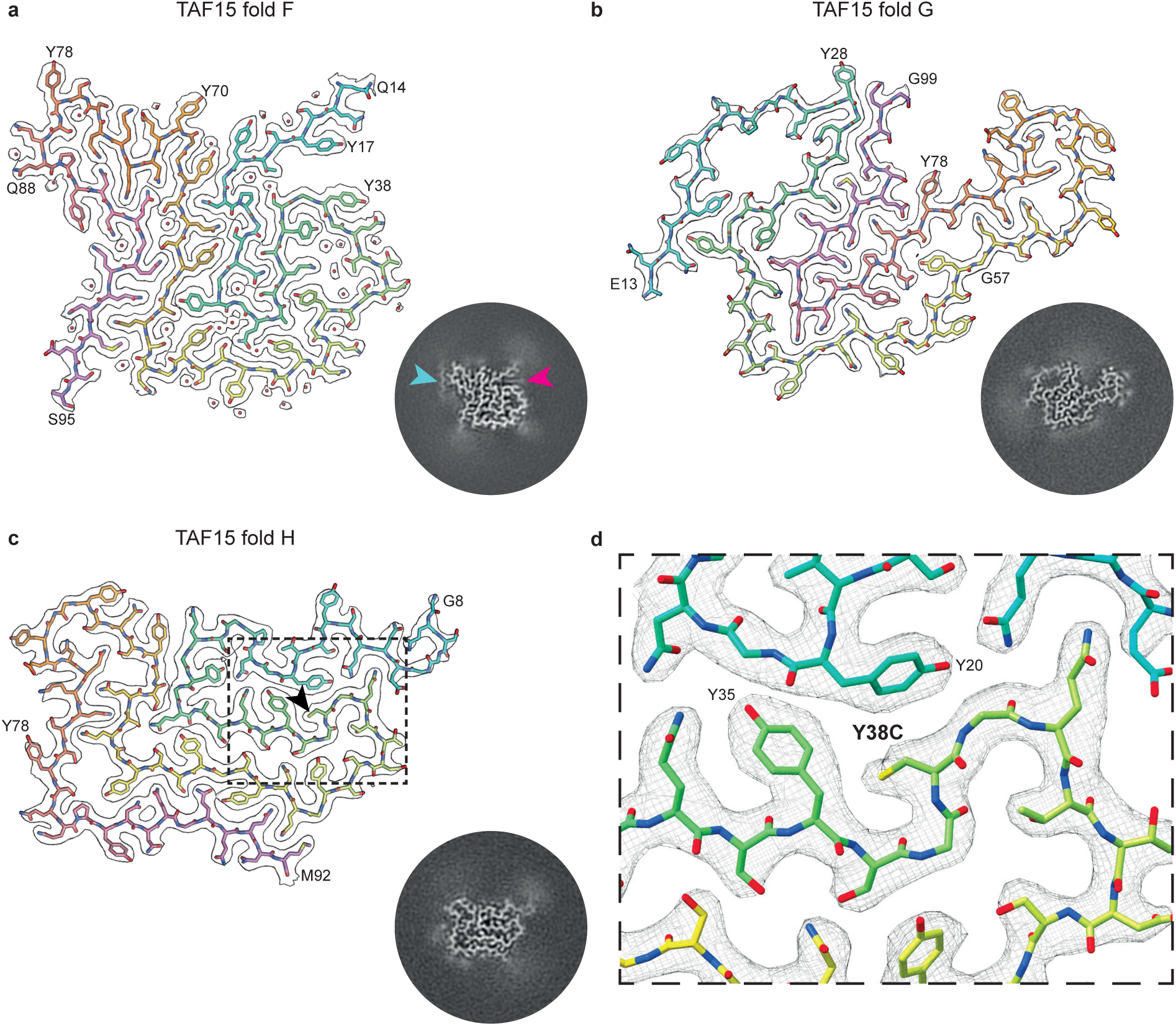
TAF15 filament folds F, G and H in BIBD. a-c, Cryo-EM reconstructions (black outlines) and atomic models of TAF15 filament fold F **(a)**, fold G **(b)** and fold H **(c)**, shown for a single TAF15 molecule perpendicular to the helical axis. The cryo-EM reconstructions are shown using a surface zone around the atomic models. The atomic models are colour-coded as a gradient from N– to C-terminal residues. The Y38C substitution in TAF15 fold H is indicated with a black arrow. Central slices of the cryo-EM reconstructions centred on the helical axis are shown as inserts. A disconnected peptide (cyan arrow) and a lipid-like density (magenta arrow) associated with fold F are highlighted. **d,** Insert of TAF15 filament fold H centred on the Y38C substitution.

All three individuals with aFTLD-U (aFTLD-U cases 1-3) were characterised by the same TAF15 filament fold that we previously reported from four other individuals with aFTLD-U^3^, which we refer to as TAF15 fold A (Supplementary Figure 3 and Supplementary Table 3). This supports our previous conclusion that aFTLD-U is characterised by this specific TAF15 filament fold^3^.

### A new, common TAF15 filament fold in NIFID

Of the 11 individuals with NIFID, eight (NIFID cases 1-8) were characterised by a new TAF15 filament fold comprising residues G5-E93, which we refer to as TAF15 fold B (Figure 1, Supplementary Figure 4 and Supplementary Table 2). Viewed perpendicular to the helical axis, the fold has three meandering layers composed of G5-Y46, G47-Y67 and G68-E93, respectively. We identified three variants of this fold (variants I-III) (Figure 1), which co-existed within individual filaments and which were present at varying proportions among the cases (Supplementary Figure 4a and n). In variants I and II, there is a prominent cavity at the turn between the first two layers enclosed by residues Y28-Y53. In variant III, this cavity is collapsed, accompanied by interior-exterior inversions of segments Q30-Q33 and Q40-S45. In addition, variants II and III differ from variant I by side chain flips of Q14 and Q15.

Variant I of TAF15 fold B associated with a disconnected 10-residue peptide adjacent to Y78-Q89 in the third layer at the C-terminal end of the fold (Figure 1a). This peptide is not present in variants II and III (Figure 1a and b). To identify the peptide, we collected a large, high-magnification cryo-EM dataset, which yielded a 1.6 Å-resolution structure (Figure 1a, Supplementary Figure 3 and Supplementary Table 3). This unambiguously revealed an SYSQ motif in the peptide engaging in zipper packing with Q80 in the filament fold. This, together with other densities for large side chains, identified the peptide as Y135-N145 of TAF15 itself. The location of the peptide at the C-terminal end of the fold suggests that it may be derived from the same TAF15 molecules that form the fold, with the intervening residues S94-N134 forming part of the filament fuzzy coat.

Variants I and II of TAF15 fold B contain additional densities resembling the round polar headgroups and thin elongated acyl chains of lipids (Figure 1a and b). The acyl chain-like densities sit adjacent to the essentially hydrophobic environment between the main chains of successive TAF15 molecules, within a groove formed by Y20-S32 and on a planar surface formed by S36-Y38. These regions are not present in variant III, due to the partial collapse of the cavity formed by Y28-Y53, and no lipid-like densities were observed (Figure 1c). The headgroup-like densities appear to be associated with a larger less well-resolved halo-like density, which might correspond to a lipid micelle, as observed for filaments of α-synuclein and amyloid-β 1-40 assembled *in vitro* in the presence of lipids^25,26^. In addition to these lipid-like densities, a large round disconnected density adjacent to Y70 is apparent in the 3D reconstructions of variants I and III (Figure 1a and c).

### Additional new TAF15 filament folds in NIFID

Unexpectedly, the remaining three individuals with NIFID (NIFID cases 9-11) were each characterised by distinct TAF15 filament folds, which we refer to as TAF15 folds C, D and E (Supplementary Table 2). TAF15 fold C, which characterised NIFID case 9, comprises TAF15 residues Y20-Q98 and exhibits the antiparallel packing of an N-terminal segment formed of Y20-S61 against the C-terminal segment of G62-Q98 (Figure 2a and Supplementary Figure 5). This C-terminal segment is creased at two places, resulting in an S-shaped six-layered fold. We also found rare C2-symmetric anti-parallel dimers of filaments with variant I of fold C, which comprised ∼7% of the filament segments (Supplementary Figure 5). The filament interface is stabilised by hydrogen bonds between the hydroxyl and peptide groups of S36 and the hydroxyl group of Y38 from both filaments, in addition to hydrophobic interactions between the aromatic side chains of Y38 (Supplementary Figure 5n). A minor variant II of fold C co-existed within individual filaments, comprising ∼2% of the filament segments, which has an N-terminal extension from D3 to T19 that forms a seventh layer adjacent to residues Y20-S32 (Supplementary Figure 5o).

TAF15 fold D, which characterised NIFID case 10, comprises TAF15 residues S6-G99 (Figure 2b and Supplementary Figure 6). This fold consists of a four-layered body formed by residues S6-G57 and N91-G99, plus a two-layered extended arm formed by residues Q58-Q90. The structure of the arm is stabilised by zipper packing of five glutamine residues (Q58, Q60, Q86, Q88 and Q90), the side chains of which form hydrogen bonds with peptide groups in the opposite layers, as well as within ladders along the filament axis. A disconnected approximately 10-residue peptide-like density sits adjacent to Q75-Y83 in the extended arm and may form zipper packing with Q81. We also found a rare population of filaments with a related fold in NIFID case 10, comprising ∼1% of the filament segments, which we refer to as TAF15 fold D’ (Figure 2c and Supplementary Figure 6). It has a similar, but not identical, body composed of residues S6-G57 and N91-G100, and an arm formed by residues Q58-Q90 (Supplementary Figure 6j). However, unlike fold D, the arm is bent into a new four-layered arrangement and touches the body in fold D’. This leads to a more compact fold and the loss of the disconnected peptide. In addition to their similar substructure, folds D and D’ share a large round disconnected density adjacent to Y46 (Figures 2b and c). We do not refer to fold D’ as a variant of fold D because we did not find evidence for their co-existence in individual filaments, which distinguishes NIFID case 10 from the other cases of NIFID.

TAF15 fold E, which characterised NIFID case 11, comprises TAF15 residues S4-G100 (Figure 2d and Supplementary Figure 7). This fold can be divided into two lobes, a large N-terminal lobe in which residues S4-G68 form four meandering layers and a smaller C-terminal lobe consisting of a hairpin arrangement of residues G69-G100. The two lobes are separated by a small solvent-filled cavity. We identified a minor co-existing second variant of this fold within filaments, comprising ∼1% of the filament segments, in which the peptide bond of G68 is flipped and the lobes are slightly repositioned (Supplementary Figure 7h). This is accompanied by the expulsion of a water molecule hydrogen-bonded to the side chains of S64 and Y70 in variant I, which become directly hydrogen-bonded in variant II. This suggests dynamic hinging of the two lobes around G68.

### Multiple new TAF15 filament folds in BIBD

The three individuals with BIBD (BIBD cases 1-3) were each characterised by distinct TAF15 folds, which we refer to as TAF15 folds F, G and H (Supplementary Table 2). TAF15 fold F, which characterised BIBD case 1, comprises residues Q14-S95 arranged in five layers (Figure 3a and Supplementary Figure 8). The first four layers, formed of Q14-G69, have a Greek key-like arrangement, with the first two layers enclosed between the third and fourth layers. A short hairpin formed of Y70-G87 protrudes perpendicular to the layers at the beginning of the fifth layer, with the C-terminal residues Q88-S95 of this layer packing against the fourth. A disconnected peptide-like density of approximately 9-residues in length sits adjacent to Y78-Q81 at the hairpin turn and may form zipper packing with Q80 and Q81. An additional lipid acyl chain-like density sits adjacent to the essentially hydrophobic environment between TAF15 rungs within a groove formed by Y17-T19 and S36-Y38.

TAF15 fold G, which characterised BIBD case 2, comprises residues E13-G99, also in a five-layered arrangement (Figure 3b and Supplementary Figure 8). Layers two to five, formed of Y28-G99, have a different Greek key-like arrangement in which layers four and five are enclosed between layers two and three, with a boot-shaped hairpin formed of Y56-Y78 extending from the compact part of the fold. The first layer is composed of the N-terminal residues E13-G27, which loops back on the Greek key-like arrangement to enclose a large cavity.

TAF15 fold H, which characterised BIBD case 3, comprises residues G8-M92 (Figure 3c and Supplementary Figure 8). The fold consists of four meandering layers, with a boot-shaped hairpin formed of Y56-Y78 between layers three and four folding back perpendicular to the other layers.

### Cryo-EM identification of a *TAF15* missense variant in BIBD

Against our expectations, the high-resolution (2.7 Å) cryo-EM reconstruction of TAF15 fold H from BIBD case 3 did not contain density for the large side chain of Y38, despite being in the interior of the ordered filament core (Figure 3d and Supplementary Figure 8). Moreover, a tyrosine at this position could not be accommodated in the fold due to steric hinderance from the adjacent Y20 and Y35 residues. Therefore, we hypothesised that this individual might have genetic variation in their *TAF15* gene resulting in an amino acid with a smaller side chain at position 38 that would fit the density and stereochemical environment of the fold. Indeed, sequencing of this individual’s *TAF15* gene revealed a heterozygous c.113A>G, p.Y38C, single nucleotide variant, whereas all other individuals had wild-type *TAF15* (Supplementary Table 1). This is an extremely rare variant, with a total allele frequency of 6.8 × 10^−6^ in the Genome Aggregation Database (GnomAD, v4.1.0). The individual had no known positive family history for neurodegenerative diseases and genetic analysis of family members was not possible. However, the fact that the filaments were formed entirely of mutated Y38C TAF15, despite the variant being heterozygous, suggests that this amino acid substitution drives filament formation and may be pathogenic, analogous to pathogenic missense single nucleotide variants that enhance the amyloid filament formation of other proteins in neurodegenerative diseases, such as tau, α-synuclein and TDP-43 (Ref ^1^).

### Shared features of TAF15 filament folds

In this work, we determined a total of 32 TAF15 filament structures from 17 individuals, comprising nine distinct TAF15 filament folds and four fold variants (Figure 4a). This rich collection of structures enabled us to identify shared features of TAF15 filament folds. The conserved sequences that form the ordered cores of the TAF15 filaments contain significantly fewer hydrophobic and aliphatic (alanine, valine, isoleucine and leucine) residues than the sequences of other amyloid filament cores, including those of tau, α-synuclein, TDP-43, amyloid-β and TMEM106B from human brain^2^ (Figure 4b). As such, TAF15 filament folds are less stabilised by hydrophobic interactions than other amyloid filament folds. Ionic interactions of amino acid side chains are also rare, with only 6% of total residues having charged side chains. Instead, our high-resolution structures show that TAF15 folds are biased towards stabilisation by extensive hydrogen bonding networks involving buried water molecules, neutral polar amino acid side chains (serine, threonine, asparagine, glutamine and tyrosine; 69% of total residues) and main chain peptide groups (especially glycine; 20% of total residues). Zipper packing of the abundant glutamine and asparagine residues (26% of total residues) is also prominent.

**Figure 4:**
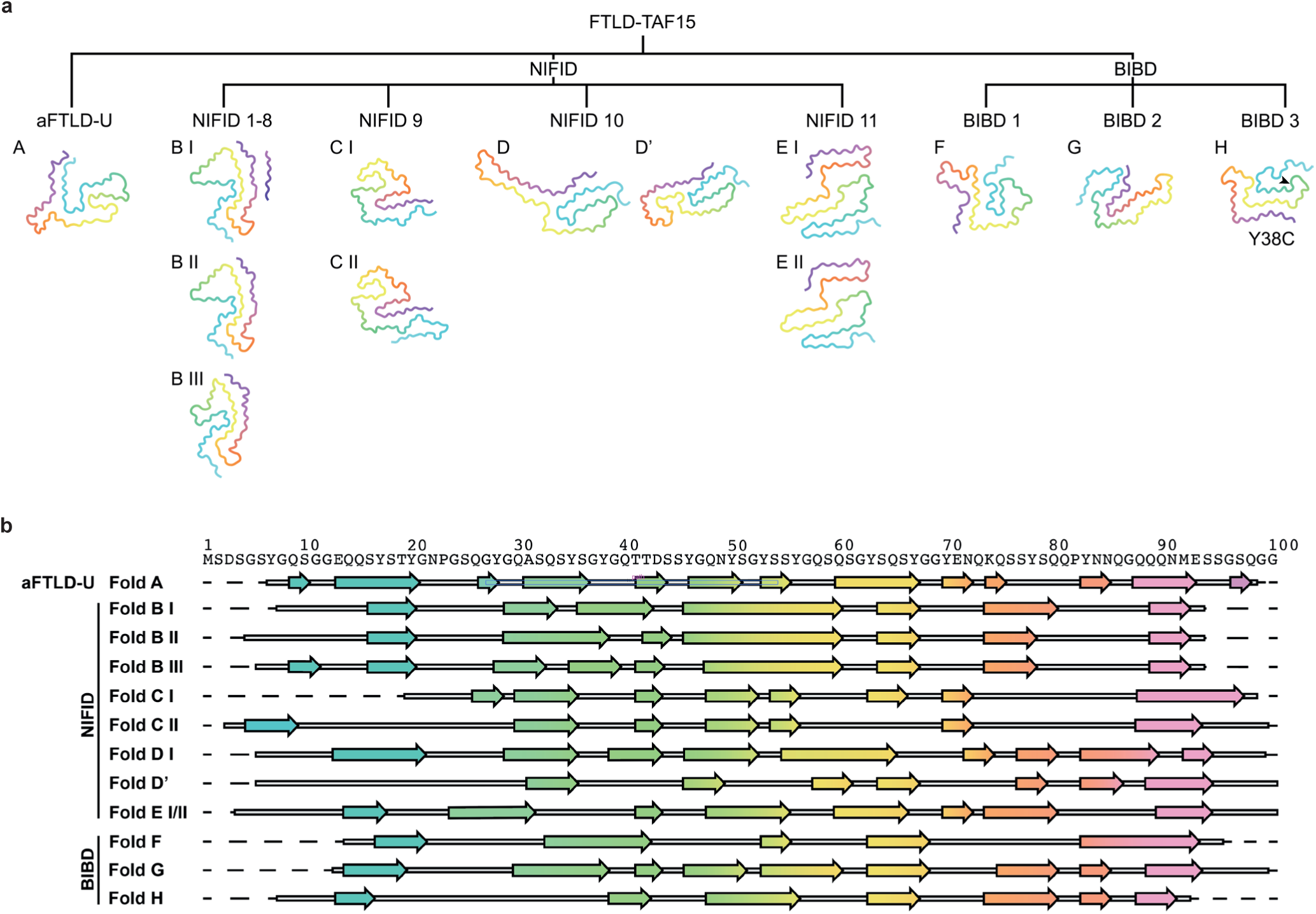
Comparison of TAF15 filament folds in aFTLD-U, NIFID and BIBD. a, Main chain traces of TAF15 filament folds (A-H) and their variants (I-III) from aFTLD-U, NIFID (cases 1-11) and BIBD (cases 1-3). The Y38C substitution in TAF15 fold H from BIBD case 3 is highlighted. Residues are colour-coded as a gradient from N– to C-terminal. **b,** Amino acid sequence alignment of the secondary structure elements of the TAF15 filament folds (A-H) and their variants (I-III). Arrows indicate β-strands, which are colour-coded as a gradient from N– to C-terminal.

Stacking of TAF15 molecules within filaments is stabilised by intermolecular β-sheets; hydrogen bonding ladders among the amide side chains of glutamine and asparagine residues; and stacking interactions of the aromatic side chains of tyrosine residues. These interactions are common among amyloid filaments^7^. The latter two interactions are enriched in TAF15 filaments due to the abundance of glutamine (6% of total residues), asparagine (20% of total residues) and tyrosine (15% of total residues) residues (Figure 4b).

We identified several recurrent hairpin-like motifs that were shared among TAF15 folds, which are stabilised by internal interactions and may represent autonomous folding units. A boot-shaped hairpin formed by Y56-Y78, exhibiting zipper packing among glutamine residues 58, 60 and 75, is the largest of these recurrent motifs and common to folds B, G and H (Supplementary Figure 9a). A smaller hairpin formed by E71-Q86, in which Q73, N84 and Q86 form zipper packing, is shared by folds A and F (Supplementary Figure 9b). Both hairpin motifs contain an exterior salt bridge between E71 and K74. This salt bridge, involving the only positively charged residue in the fold-forming sequence, is present in all folds except fold D, in which E71 is buried in the fold interior. Folds C and G share a hairpin formed by Y78-S95, exhibiting zipper packing between Q80 and E93, and between Q84 and Q89 (Supplementary Figure 9c). Folds A and E feature a hairpin formed by Y53-Y63 in which Y53 and S61 interact (Supplementary Figure 9d).

Several of the TAF15 folds were associated with similar additional densities. Disconnected peptide-like densities of 9-10 residues in length were found adjacent to the region Y75-Q89 in variant I of fold B and in folds D and F (Figures 1a, 2b and 3a). For variant I of fold B, our high-resolution reconstruction revealed an SYSQ motif present in the peptide, identifying it as Y135-N145 of TAF15 itself (Figure 1a). No unambiguous sequence motifs were identified in the peptides associated with the other two folds, but the similar positions of the disconnected peptides near the C-terminal ends of the respective folds and apparent zipper packing with Q80 and/or Q81 of the respective TAF15 folds suggest that these peptides could also be similar ordered segments of the C-terminal fuzzy coat.

Extended lipid-like densities were associated with a shared tyrosine arc ^35^YSGYG^39^ motif in variants I and II of TAF15 fold B and in fold F (Figures 1a and b and 3a), but not with variant III of fold B or with the other folds. Their selective association suggests these densities may represent endogenous lipids bound to particular types of TAF15 filaments. However, the composition of such lipids is most likely altered by the use of detergent during filament extraction, thus hindering the elucidation of their chemical identity. Large round disconnected densities in proximity to the exterior-facing side chains of Y46 or Y70 were associated with variants I and II of fold B and with folds D and D’ (Figures 1a and b and 2b and c). Attempts to improve the resolution of these densities using focussed refinements and alternative symmetries were unsuccessful, possibly because the densities comprise a mixture of macromolecules. Future work will be required to identify these different densities, as well as to assess if they act as cofactors for filament formation and if they have any functional consequences for neurons.

### Cryo-EM-guided reappraisal of the classification of FTLD-FET subtypes

While a single TAF15 fold characterised all cases of aFTLD-U (fold A), four TAF15 folds were associated with NIFID (folds B-E) and three folds were associated with BIBD (folds F-H) (Figure 4a). Of the folds associated with NIFID, fold B was the most common, characterising eight of the 11 cases. FET protein-immunoreactive NCIs in the prefrontal cortex in aFTLD-U are uniformly compact round, whereas more heterogeneous types of inclusions are associated with NIFID and BIBD, including compact round, annular and tangle-like NCIs, as well as dystrophic neurites and neuropil dots^18,23^. We hypothesised that the different inclusion types in NIFID and BIBD might segregate with the distinct TAF15 folds found using cryo-EM. To test this, we undertook a blinded reappraisal and semi-quantitative assessment of TAF15 cytoplasmic inclusion morphology in prefrontal cortex sections from the 17 individuals with FTLD-FET that we analysed using cryo-EM (Figure 5 and Supplementary Table 2).

**Figure 5:**
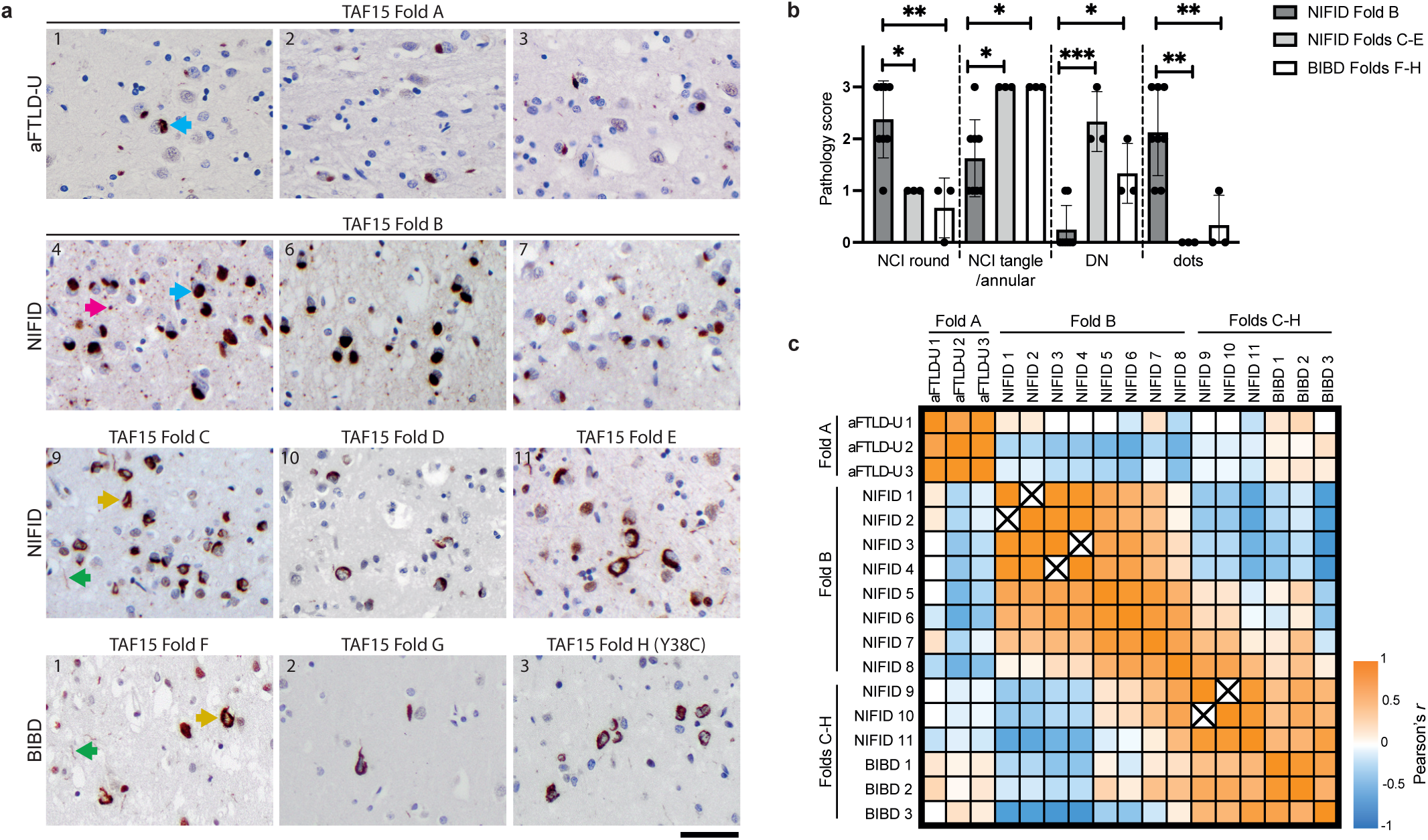
Relationship between TAF15 filament folds and cytoplasmic inclusions. a, Immunohistochemistry for TAF15 (brown) in prefrontal cortex sections from aFTLD-U cases 1-3 with TAF15 filament fold A; NIFID cases 4, 6 and 7 with TAF15 fold B; NIFID cases 9-11 with TAF15 folds C-E, respectively; and BIBD cases 1-3 with TAF15 folds F-H, respectively. Sections were counterstained with haematoxylin (blue). The predominant TAF15-immunoreactive neuronal cytoplasmic inclusions (NCIs) in aFTLD-U cases are compact round (cyan arrows). NIFID cases with fold B are characterised by a predominance of compact round NCIs and the presence of neuropil dots (magenta arrow). NIFID cases with folds C-E and BIBD cases with folds F-H show a predominance of annular and tangle-like NCIs (yellow arrows) and the presence of dystrophic neurites (DN, green arrows). Case numbers are indicated at the top left of the images. Scale bar, 25 µm. **b,** Graph of semi-quantitative scoring of TAF15-immunoreactive cytoplasmic inclusion types among NIFID cases with fold B (n=8), NIFID cases with folds C-E (n=3) and BIBD cases with folds F-H (n=3) from Supplementary Table 3. Individual datapoints and means ± SD are shown. *p<0.05, **p<0.01, ***p<0.001, (Kruskal-Wallis test with Dunn’s post-hoc test). **c,** Heatmap of correlation (Pearson’s *r*) among semi-quantitative scoring of TAF15-immunoreactive cytoplasmic inclusion types in the aFTLD-U (n=3), NIFID (n=11) and BIBD (n=3) cases from Supplementary Table 3. Crosses indicate that the values for both variables are identical, precluding correlation analysis. The TAF15 filament folds (A-H) of the cases are indicated.

The cases of aFTLD-U were all characterised by uniform compact round inclusions, consistent with previous reports^4,18^. Notably, the inclusion types were also clearly delineated between the eight individuals with NIFID that shared TAF15 fold B (cases 1-8) and the remaining three individuals each with NIFID (cases 9-11) and BIBD (cases 1-3), which each had distinct TAF15 filament folds (folds C-H) (Figures 5a and b). The NIFID cases with fold B were characterised by a predominance of compact round NCIs and the presence of neuropil dots, with few annular and tangle-like NCIs or neuritic inclusions. By contrast, the remaining NIFID cases and the BIBD cases were characterised by a predominance of more variable inclusion morphologies, including annular and tangle-like NCIs, and dystrophic neurites, with few compact round NCIs and neuropil dots. This shows that the NIFID cases with TAF15 fold B are structurally and neuropathologically distinct compared to the other cases of FTLD-FET examined (Figure 5c).

The remaining three cases of NIFID and the three cases of BIBD could not be distinguished from one another based on the neuropathological re-assessment of NCI morphology and distribution (Figure 5 and Supplementary Table 2). Conceivably, this may be due to a lack of statistical power because we only identified one case each for TAF15 filament folds C-H. Assessment of inclusion types in cases with these folds should be revisited as frozen tissue from additional cases of NIFID and BIBD becomes available in the future.

## DISCUSSION

We previously showed that TAF15 amyloid filaments characterised all four examined cases of the FTLD-FET subtype aFTLD-U^3^, thereby adding TAF15 to the small group of amyloid filament-forming proteins that characterise neurodegenerative diseases^2^. However, it was unclear if TAF15 amyloid filaments were more widespread in neurodegenerative diseases, especially in the two additional proposed subtypes of FTLD-FET, NIFID and BIBD, which are associated with diverse clinical presentations including FTD, MND and movement disorders^5,6,18^.

Here, we determined a total of 32 amyloid filament structures from 17 cases of FTLD-FET, comprising 3 additional cases of aFTLD-U, 11 cases of NIFID and 3 cases of BIBD, and encompassing the diverse clinical presentations of this group of diseases. Against previous assumptions, all cases were characterised by TAF15 amyloid filaments. This provides strong evidence that TAF15 amyloid filaments define all three proposed subtypes of FTLD-FET and establishes a group of TAF15 proteinopathies associated with FTD, MND and movement disorders. This is analogous to tauopathies, synucleinopathies and TDP-43 proteinopathies, whereby intracellular amyloid filaments of these proteins in the CNS each characterise multiple neurodegenerative diseases^2,27^. As such, we propose that the term FTLD-TAF15 should now replace both FTLD-FET and its predecessor FTLD-FUS.

As in our previous cryo-EM study of four aFTLD-U cases^3^, we did not detect any filaments composed of the other two FET proteins, FUS and EWS, in the 17 cases of FTLD-FET analysed here. This strongly suggests that FUS and EWS do not form amyloid filaments in FTLD-FET. We hypothesise that the immunoreactivity of inclusions for FUS and EWS, as well as for the FET protein nuclear import receptor transportin-1^28,29^, in FTLD-FET most likely represents the sequestration of non-filamentous species of these proteins by TAF15 filaments, either directly or through shared binding partners^13^. It remains possible that FUS filament formation plays a role in cases of ALS caused by pathogenic mutations in *FUS*^16,17^, in which NCIs are immunoreactive against FUS, but not TAF15, EWS or transportin-1 (Refs^14,29^). This may be analogous to the filamentous inclusions of superoxide dismutase 1 that characterise cases of ALS with pathogenic *SOD1* mutations^30^. Cryo-EM structures of putative amyloid filaments from these genetic forms of ALS are awaited to test this hypothesis.

Amyloid filament formation of TAF15 and not of FUS or EWS is supported by the strong nuclear depletion of TAF15 in neurons containing FET protein-immunoreactive NCIs, whereas FUS and EWS are only mildly depleted^14^. This may be analogous to the nuclear depletion of TDP-43 upon the formation of intracellular TDP-43 filament inclusions^31–33^. For TDP-43, nuclear depletion results in multiple loss of function effects^34^, which may contribute to neurodegeneration. Future studies should investigate if nuclear depletion of TAF15 as a consequence of its filament formation results in disease-relevant loss of function effects^35^.

We found a total of seven distinct TAF15 filament folds among 17 cases of FTLD-FET. All of the folds were formed by essentially the same N-terminal 100 residues of TAF15. Two isoforms of TAF15 have been reported, which differ by the inclusion or exclusion of ^60^QSG^62^ in the LCD following alternative splicing of exon 4 (Refs^36–38^). All of the TAF15 folds contained residues ^60^QSG^62^, establishing that the filaments were formed exclusively of the longer reported isoform. If the shorter TAF15 isoform exists in the CNS, it is possible that it might protect against filament formation. This would be distinct from the alternatively spliced CNS isoforms of tau, which form filaments with distinct core folds in different diseases^39^. Future studies should assess the expression of TAF15 isoforms in the human CNS.

Unexpectedly, we found four distinct TAF15 filament folds among the 11 cases of NIFID. While eight NIFID cases shared a fold (fold B), the remaining three cases each had a different fold (folds C, D and E). Furthermore, each of the three BIBD cases were characterised by distinct TAF15 folds (fold F, G and H). This was not the case for the three cases of aFTLD-U, which were all characterised by the same TAF15 fold that we previously found in four separate cases of aFTLD-U (fold A)^3^. This suggests that, while aFTLD-U is a single disease entity, NIFID and BIBD may each comprise multiple disease entities, resulting in a much greater number of FTLD-FET subtypes than previously thought.

The homogeneous compact round morphology of TAF15 NCIs in aFTLD-U clearly distinguishes it from the other FTLD-FET subtypes, supporting the classification of aFTLD-U as a single disease entity. Neuropathological re-evaluation of neocortical TAF15 inclusion morphology established that the eight NIFID cases with the common TAF15 fold B exhibited a unifying neocortical TAF15 inclusion pathology, which was distinct from the other cases of FTLD-FET examined. This provides additional evidence that the NIFID cases with TAF15 fold B represent a distinct disease entity. This is similar to the previous identification of multiple tau filament folds in progressive supranuclear palsy (PSP) using cryo-EM, which also aligned with neuropathological re-evaluation and supported a new disease entity, limbic-predominant neuronal inclusion body 4R tauopathy (LNT)^39^. Together with our results, this shows that cryo-EM can expand neuropathology and the molecular definition of diseases.

We did not observe pronounced differences in the more heterogeneous neocortical TAF15 inclusion morphologies among the three cases of NIFID and the three cases of BIBD that each had a distinct TAF15 fold. Neuropathological correlation was limited for these folds since they were only identified in a single case each and should be revisited as additional cases of NIFID and BIBD become available. As such, it is currently less clear if these TAF15 folds are each associated with distinct disease entities, or if multiple folds might converge on single disease entities. The latter has not previously been observed for amyloid filament folds of other neurodegenerative disease-associated proteins^2^. We anticipate that neuropathological and cryo-EM analyses of additional cases of NIFID and BIBD, when available, will enable the re-classification of additional subtypes of FTLD-TAF15, analogous to the different types of FTLD-TDP^15^, with the presence of intermediate filament-immunoreactive inclusions and basophilic inclusions being secondary neuropathological features.

Unlike the homogenous bvFTD clinical presentation of aFTLD-U, the NIFID and BIBD cases had heterogenous clinical presentations of FTD, MND and/or extrapyramidal dysfunction, which did not correlate with TAF15 filament folds. There were also no significant differences in age at onset and disease duration among the 17 cases, although we note that the aFTLD-U and the NIFID cases with fold B had a trend towards younger age at onset than the other cases. Additional studies of larger cohorts, when available, will be required to test if there are significant differences in clinical presentation among NIFID and BIBD cases with distinct TAF15 filament folds and inclusion pathologies. We note that overlapping clinical presentations are not uncommon among distinct neurodegenerative diseases, for example among different subtypes of FTLD-TDP and tauopathies^15^.

The mechanisms underlying the formation of distinct filament folds in different neurodegenerative diseases and their consequences are not understood. They may relate to the formation of filaments in distinct cellular environments in different diseases, followed by templated seeded assembly during prion-like propagation^2,40^. The filament folds identified here will guide the development of model systems and investigation of the molecular mechanisms underlying TAF15 proteinopathies. They will also aid in the development of fold-specific binders, which have utility for disease diagnosis and monitoring^41–44^, as well as for the inhibition of filament formation and disassembly of filaments as potential therapeutic strategies^45,46^. Currently, there are no diagnostic or therapeutic tools for FTLD-FET.

The high-resolution cryo-EM structure of filaments with TAF15 fold H from BIBD case 3 showed that they were composed entirely of TAF15 with a Y38C amino acid substitution. This led to the identification that this individual had an extremely rare heterozygous c.113A>G (p.Y38C) single nucleotide variant in *TAF15*. Given that the filaments were composed entirely of Y38C TAF15, despite the individual being heterozygous, we propose that this variant drives the assembly of TAF15 into amyloid filaments. We, therefore, speculate that this mutation may be pathogenic, analogous to pathogenic missense variants in other proteins including tau, α-synuclein and TDP-43 that enhance their amyloid filament formation^1^. Less-rare variants in *TAF15*, which mostly reside outside of the LCD, have previously been detected in ALS patients in the absence of positive family histories^47–49^. Unlike in our study, it was not clear if these patients had TAF15 pathology. Genetic variation associated with FTLD-FET has so far remained elusive, likely because of the historical focus on *FUS* rather than *TAF15*.

## CONCLUSION

This study establishes that all three proposed subtypes of FTLD-FET – aFTLD-U, NIFID and BIBD – are characterised by TAF15 amyloid filaments, motivating their reclassification as FTLD-TAF15. A distinct TAF15 filament fold and inclusion morphology characterised aFTLD-U, demonstrating that this is a distinct disease entity. In contrast, NIFID and BIBD were associated with multiple TAF15 filament folds and inclusion morphologies, suggesting that they may each comprise multiple disease entities. This was supported by neuropathological reassessment of their TAF15 filament inclusion morphologies. Our identification of a heterozygous *TAF15* variant that led to filaments composed entirely of the mutated protein provides structural and genetic evidence for TAF15 proteinopathy in neurodegenerative diseases. This structure-based classification of TAF15 proteinopathies will guide research into their underlying disease mechanisms, as well as the development of diagnostic and therapeutic tools.

## METHODS

### Statistical analyses

A Kruskal-Wallis test with Dunn’s post-hoc test was used to determine if there were significant differences between the semi-quantitative scoring of TAF15-immunoreactive neuronal cytoplasmic inclusion types among NIFID cases with TAF15 filament fold B (n=8), NIFID cases with other TAF15 filament folds (n=3) and BIBD cases (n=3). Pearson correlation was used to determine the correlation among the semi-quantitative scoring of TAF15-immunoreactive neuronal cytoplasmic inclusion types in the aFTLD-U, NIFID and BIBD cases. Statistical analyses were performed in GraphPad Prism (version 10.4.2).

### Human tissue samples

Postmortem CNS tissue samples from 17 individuals with a neuropathological diagnosis of FTLD-FET were obtained from the DZNE Brain Bank; the University of British Columbia Brain Bank; the University of Alberta; the University of Calgary; Western University, London, Canada; the Queen Square Brain Bank for Neurological Disorders at Queen Square Institute of Neurology, University College London; and the Dementia Laboratory Brain Library at Indiana University School of Medicine. Consent for autopsy was obtained from the legal representative in accordance with local institutional review boards.

### Neuropathological analysis

Cases were selected based on a neuropathological diagnosis of aFTLD-U (n=3), NIFID (n=11) or BIBD (n=3). Neuropathological diagnosis was performed according to^50^. Atypical FTLD-U was diagnosed by the presence of FET protein-immunoreactive NCIs, in the absence of NCIs visible by H&E and α-internexin-immunoreactive NCIs. NIFID was diagnosed by the presence of FET protein-immunoreactive NCIs, in addition to the presence of round pick-body-like NCIs and/or hyaline conglomerate inclusions visible by H&E (usually present in most cortical regions including the prefrontal cortex), and α-internexin-positive inclusions. BIBD was diagnosed by the presence of FET protein-immunoreactive NCIs and basophilic NCIs visible by H&E (usually present in basal ganglia, midbrain and spinal cord, and rare or absent in the prefrontal cortex), together with the absence of α-internexin-positive inclusions. Demographic, clinical and genetic data are given in Supplementary Table 1. Pathological details are given in Supplementary Table 2.

Re-evaluation of the neuropathological diagnosis and semi-quantitative assessment of TAF15-immunoreactive inclusion types was performed in a blinded and centralised manner. The prefrontal cortex, the brain region used for cryo-EM analyses, was investigated for all cases. For selected cases, additional regions including hippocampus, basal ganglia and spinal cord were included. Semi-quantitative assessment used the following grading scheme: 0, absent; 1, mild/rare (only a few examples in the examined section); 2, moderate (readily observable but not in every imaging field); 3, numerous/frequent (readily observable in every imaging field). For evaluation of the morphological subtype of NCIs (compact round, annular and tangle-like), the following grading scheme was used: 0, absent; 1, <25% of NCIs; 2, between 25% and 75% of NCIs; 3, >75% of NCIs..

### Genetic analysis

Short-read genome sequencing data for all cases except NIFID cases 2, 3 and 6-10 were generated using the Nextera DNA Flex Library prep kit, followed by sequencing on an Illumina NovaSeq. Reads were mapped to the human reference sequence (GRCh38 build) using BWA-MEM^51^, followed by variant calling using GATK HaplotypeCaller according to the GATK best practices^52^. *TAF15* variants were subsetted and normalized using BCFtools^53^, then annotated with the Ensembl Variant Effect Predictor^54^. For NIFID cases 2, 3 and 6-10, the 16 coding exons of *TAF15* were amplified using polymerase chain reactions and analyzed by Sanger sequencing.

### Extraction of sarkosyl-insoluble proteins from human brain

Sarkosyl-insoluble proteins were extracted from flash-frozen prefrontal cortex samples and stored at –70°C as previously described^3^. Grey matter was dissected and homogenized using a Polytron (Kinematica) in 40 volumes (v/w) of extraction buffer (10 mM Tris-HCl pH 7.4, 0.8 M NaCl, 10% sucrose, 1 mM dithiothreitol and 1 mM EGTA) containing protease and phosphatase inhibitor cocktail (Pierce). A 25% solution of sarkosyl in water was added to the homogenates to yield a total concentration of 2% sarkosyl. The homogenates were then incubated for 1 h or overnight at 37 °C with orbital shaking at 200 RPM, followed by centrifugation at 27,000 *g* for 10 min. The supernatants were divided into 1 mL aliquots and centrifuged at 166,000 *g* for 20 min. Each pellet was soaked in 20 µL extraction buffer containing 1% sarkosyl at 37 °C for ≥20 min, then resuspended by extensive pipetting. Aliquots were combined in batches of six, topped up to 0.5 mL with extraction buffer and sonicated for 5 min at 50% amplitude (Qsonica Q700). The samples were then diluted to 1 mL with the same buffer and centrifuged at 17,000 *g* for 5 min to remove any large particulates. The supernatants were retained and centrifuged at 166,000 *g* for 20 min. The resulting pellets were soaked and resuspended as before then combined. The sample was topped up to 1 mL with extraction buffer containing 1% sarkosyl and incubated overnight at 37 °C with orbital shaking at 200 RPM. The samples were centrifuged at 166,000 *g* for 20 min and the pellets were resuspended in 25 µL/g tissue of 20 mM Tris-HCl pH 7.4, 150 mM NaCl by soaking at 37 °C, extensive pipetting and sonication for 5 min at 50% amplitude. All centrifugation steps were carried out at 25 °C.

### Immunolabelling

#### Immunohistochemistry

Immunohistochemistry was performed on 5 µm-thick paraffin-embedded, formalin-fixed tissue sections using antibodies against FUS (Proteintech 60160-1-Ig) at a dilution of 1:2,000; TAF15 (Bethyl IHC-00094) at a dilution of 1:300; and α-internexin (Invitrogen 32-3600) at a dilution of 1:1,000 using a Ventana BenchMark GX automated staining system (Roche) using the optiVIEW DAB detection kit (Roche). Heat-induced antigen retrieval was performed for all antibodies with Cell Conditioning 1 (CC1) buffer (Roche).

#### Immunoblotting

For immunoblotting, sarkosyl-insoluble extracts from 0.1 g tissue were diluted 30-fold in lithium dodecyl sulfate sample buffer (Thermo) containing a total concentration of 1% β-mercaptoethanol and heated at 95°C for 10 min. Samples were resolved using 4–12% BIS-Tris gels (Novex) at 200 V for 40 min and transferred onto nitrocellulose membranes. Membranes were blocked in PBS containing 1% BSA and 0.2% Tween for 30 min at 21 °C, then incubated in the same buffer for 1 h at 21 °C with primary antibodies against FUS (Proteintech 11570-1-AP) used at a dilution of 1:5,000; TAF15 (Bethyl A300-308A) used at a dilution of 1:5,000; and EWS (Santa Cruz sc-28327) used at a dilution of 1:250. Membranes were then washed three times with PBS containing 0.2% Tween and incubated with Goat Anti-Mouse IgG StarBright Blue 700 (Biorad) or Anti-Rabbit IgG DyLight 800 4x PEG Conjugate (Cell Signal Technology) secondary antibodies for 1 h at 21 °C. Membranes were then washed three times with PBS containing 0.2% Tween and imaged using a ChemiDoc MP (Bio-Rad).

### Cryo-EM

Sarkosyl-insoluble extracts from 1-2 g tissue were incubated with 0.4 mg/mL pronase (Sigma) for 1 h at 21 °C and centrifuged at 3,000 *g* for 15 s immediately before plunge-freezing to remove large particulates. Three µL of the supernatant was applied to glow-discharged 1.2/1.3 μm holey carbon-coated 200-mesh gold grids (Quantifoil) and plunge-frozen in liquid ethane using a Vitrobot Mark IV (Thermo Fisher) with a blot force of 12-14 for 4-8 s at 4 °C and 100% humidity. Images were acquired using a 300 keV Titan Krios microscope (Thermo Fisher) with either a K3 detector (Gatan) and a GIF-quantum energy filter (Gatan) operated at a slit width of 20 eV, or a Falcon 4 detector (Thermo Fisher Scientific). Aberration-free image shift (AFIS) within the EPU software (Thermo Fisher) was used during image acquisition. Further details are given in Supplementary Table 3.

### Helical reconstruction

Movie frames were gain-corrected, aligned, dose-weighted and summed using the motion correction program in RELION-5.0 (Ref ^55^). The motion-corrected micrographs were used to estimate the contrast transfer function (CTF) using CTFFIND-4.1^56^. All subsequent image-processing was performed using helical reconstruction methods in RELION-5.0^24,57^. Amyloid filaments were picked manually and reference-free two-dimensional (2D) classification was performed to remove suboptimal segments. Initial three-dimensional (3D) reference models were generated *de novo* by producing sinograms from 2D class averages as previously described^58^. Unmasked, followed by masked 3D autorefinements with optimisation of the helical twist were performed, followed by iterative Bayesian polishing and CTF refinement^55,59^. Where beneficial, 3D classification with or without alignments was used to further remove suboptimal segments. 3D autorefinement, Bayesian polishing, and CTF refinement were then repeated. The final reconstructions were sharpened using the standard post-processing procedures in RELION-5.0 and overall resolutions were estimated from Fourier shell correlations of 0.143 between the two independently refined half-maps, using phase-randomization to correct for convolution effects of a generous, soft-edged solvent mask^60^. Local resolution estimates were obtained using the same phase-randomization procedure, but with a soft spherical mask that was moved over the entire map. Helical symmetry was imposed using the RELION Helix Toolbox. Further details are given in Supplementary Table 3.

### Atomic model building and refinement

The atomic models were built *de novo* and refined in real-space in COOT^61^ using the best-resolved map for each TAF15 filament fold. Rebuilding using molecular dynamics was carried out in ISOLDE^62^. The model was refined in Fourier-space using REFMAC5^63^, with appropriate symmetry constraints defined using Servalcat^64^, or using PHENIX^65^. To confirm the absence of overfitting, the model was shaken, refined in Fourier-space against the first half-map using REFMAC5 or PHENIX and compared to the second half map. Geometry was validated using MolProbity^66^. Molecular graphics and analyses were performed in ChimeraX^61^. Secondary structure was assigned in ChimeraX using the Defining the Secondary Structure of Proteins (DSSP) algorithm^62^, with a hydrogen bond energy cutoff of –0.5 kcal/mol and a minimum strand length of two residues. Model statistics are given in Supplementary Table 3.

### Data availability

Cryo-EM datasets will be deposited to the Electron Microscopy Public Image Archive (EMPIAR) before publication. Cryo-EM reconstructions of TAF15 filaments have been deposited to the Electron Microscopy Data Bank (EMDB) under accession codes EMD-55930 (fold B variant I), EMD-55931 (fold B variant II), EMD-55932 (fold B variant III), EMD-56031 (fold C variant I singlet), EMD-56032 (fold C variant I doublet), EMD-56041 (fold C variant II), EMD-56030 (fold D), EMD-56040 (fold D’), EMD-56028 (fold E variant I), EMD-56029 (fold E variant II), EMD-56027 (fold F), EMD-56026 (fold G), EMD-56025 (fold H). Atomic models of TAF15 filaments have been deposited to the Protein Data Bank (PDB) under accession codes 9THM (fold B variant I), 9THN (fold B variant II), 9THP (fold B variant III), 9TKK (fold C variant I singlet), 9TKL (fold C variant I doublet), 9TL2 (fold C variant II), 9TKJ (fold D), 9TKZ (fold D’), 9TKH (fold E variant I), 9TKI (fold E variant II), 9TKG (fold F), 9TKF (fold G), 9TKE (fold H). Genetic sequencing data will be made publicly available before publication.

## ACKNOWLEDGEMENTS

We thank the individuals and their families for donating brain tissue; the DZNE Brain Bank, the University of British Columbia Brain Bank, the Queen Square Brain Bank for Neurological Disorders, which is supported by the Reta Lila Weston Institute of Neurological Studies, at University College London Queen Square Institute of Neurology and the Dementia Laboratory Brain Library at Indiana University School of Medicine for supplying tissue samples; staff at the MRC Laboratory of Molecular Biology Electron Microscopy Facility for access to and support with cryo-EM; staff at the MRC Laboratory of Molecular Biology Scientific Computing Facility for access to and support with computing; R. Richardson and M. Jacobsen for help with immunohistochemistry; M.R. Farlow for clinical characterisation of NIFID case 4; M. Mesulam for clinical characterisation of NIFID case 11; D. Arseni, A. Bertolotti, R. Chen, A. Giblin, M. Goedert, F. Hawkins, S. Mishra, S.H.W. Scheres and M. Walker for discussions. This work was supported by the Medical Research Council, as part of United Kingdom Research and Innovation (also known as UK Research and Innovation) (MC_UP_1201/25 to B.R.-F.); the US National Institutes of Health (R01-NS137469 to K.N. and B.R.-F; R01-NS110437, UAG063911 to I.R.A.M, P30-AG010133, RF1-AG071177 and R01-AG080001 to B.G); the Alzheimer’s Society (AS-PG-18-004 and AS-PG-21-004 to T.L.); the Association for Frontotemporal Degeneration (2019-0009 to T.L.); and a Swiss National Science Foundation Postdoctoral Fellowship (P500PB_206890 to S.T.). For the purpose of open access, the MRC Laboratory of Molecular Biology has applied a CC BY public copyright licence to any Author Accepted Manuscript version arising.

## AUTHOR CONTRIBUTIONS

S.R., J.T.J., K.N., R.C., S.D., L.-C.A., M.S., J.H., R.R., B.G., T.L., I.R.A.M. and M.N. identified individuals and performed neuropathological evaluation; M.N. performed neuropathological re-evaluation, with consensus agreement among B.G., T.L., I.R.A.M and M.N.; S.T. and N.R.V. prepared brain extracts, performed immunoblot analysis and collected cryo-EM data; S.T., N.R.V., A.G.M. and B.R.-F. analysed cryo-EM data; W.D.C., M.V.B. and R.R. performed genetic analyses; B.R.-F. supervised the study; all authors contributed to writing the manuscript.

## REFERENCES

1. Wilson, D. M. et al. Hallmarks of neurodegenerative diseases. Cell 186, 693–714 (2023).

2. Scheres, S. H. W., Ryskeldi-Falcon, B. & Goedert, M. Molecular pathology of neurodegenerative diseases by cryo-EM of amyloids. Nature 621, 701–710 (2023).

3. Tetter, S. et al. TAF15 amyloid filaments in frontotemporal lobar degeneration. Nature 625, 345–351 (2024).

4. Neumann, M. et al. A new subtype of frontotemporal lobar degeneration with FUS pathology. Brain 132, 2922–2931 (2009).

5. Neumann, M. et al. Abundant FUS-immunoreactive pathology in neuronal intermediate filament inclusion disease. Acta Neuropathol 118, 605–616 (2009).

6. Munoz, D. G. et al. FUS pathology in basophilic inclusion body disease. Acta Neuropathol 118, 617–627 (2009).

7. Sawaya, M. R., Hughes, M. P., Rodriguez, J. A., Riek, R. & Eisenberg, D. S. The expanding amyloid family: Structure, stability, function, and pathogenesis. Cell 184, 4857–4873 (2021).

8. Chothia, C. & Lesk, A. M. The relation between the divergence of sequence and structure in proteins. The EMBO Journal 5, 823–826 (1986).

9. Radamaker, L. et al. Cryo-EM reveals structural breaks in a patient-derived amyloid fibril from systemic AL amyloidosis. Nat Commun 12, 875 (2021).

10. Arseni, D. et al. TDP-43 forms amyloid filaments with a distinct fold in type A FTLD-TDP. Nature 620, 898–903 (2023).

11. Wischik, C. M. et al. Structural characterization of the core of the paired helical filament of Alzheimer disease. Proc. Natl. Acad. Sci. U.S.A. 85, 4884–4888 (1988).

12. Chiti, F. et al. Designing conditions for in vitro formation of amyloid protofilaments and fibrils. Proceedings of the National Academy of Sciences 96, 3590–3594 (1999).

13. Schwartz, J. C., Cech, T. R. & Parker, R. R. Biochemical Properties and Biological Functions of FET Proteins. Annu Rev Biochem 84, 355–379 (2015).

14. Neumann, M. et al. FET proteins TAF15 and EWS are selective markers that distinguish FTLD with FUS pathology from amyotrophic lateral sclerosis with FUS mutations. Brain 134, 2595–2609 (2011).

15. Neumann, M. & Mackenzie, I. R. A. Neuropathology of non-tau frontotemporal lobar degeneration. Neuropath Appl Neurobiol 45, 19–40 (2019).

16. Kwiatkowski, T. J. et al. Mutations in the FUS/TLS Gene on Chromosome 16 Cause Familial Amyotrophic Lateral Sclerosis. Science 323, 1205–1208 (2009).

17. Vance, C. et al. Mutations in FUS, an RNA Processing Protein, Cause Familial Amyotrophic Lateral Sclerosis Type 6. Science 323, 1208–1211 (2009).

18. Mackenzie, I. R. A. et al. Distinct pathological subtypes of FTLD-FUS. Acta Neuropathol 121, 207–218 (2011).

19. Yokoo, H., Oyama, T., Hirato, J., Sasaki, A. & Nakazato, Y. A case of Pick’s disease with unusual neuronal inclusions. Acta Neuropathol 88, 267–272 (1994).

20. Cairns, N. J. et al. Patients with a novel neurofilamentopathy: dementia with neurofilament inclusions. Neuroscience Letters 341, 177–180 (2003).

21. Nelson, J. S. & Prensky, A. L. Sporadic Juvenile Amyotrophic Lateral Sclerosis: A Clinicopathological Study of a Case With Neuronal Cytoplasmic Inclusions Containing RNA. Arch Neurol 27, 300–306 (1972).

22. Oda, M., Akagawa, N., Tabuchi, Y. & Tanabe, H. A sporadic juvenile case of the amyotrophic lateral sclerosis with neuronal intracytoplasmic inclusions. Acta Neuropathol 44, 211–216 (1978).

23. Lashley, T. et al. A comparative clinical, pathological, biochemical and genetic study of fused in sarcoma proteinopathies. Brain 134, 2548–2564 (2011).

24. He, S. & Scheres, S. H. W. Helical reconstruction in RELION. J Struct Biol 198, 163–176 (2017).

25. Frieg, B. et al. The 3D structure of lipidic fibrils of α-synuclein. Nat Commun 13, 6810 (2022).

26. Frieg, B. et al. Cryo-EM structures of lipidic fibrils of amyloid-β (1-40). Nat Commun 15, 1297 (2024).

27. Mackenzie, I. R. A. et al. Nomenclature for neuropathologic subtypes of frontotemporal lobar degeneration: consensus recommendations. Acta Neuropathol 117, 15–18 (2009).

28. Brelstaff, J. et al. Transportin1: a marker of FTLD-FUS. Acta Neuropathol 122, 591–600 (2011).

29. Neumann, M. et al. Transportin 1 accumulates specifically with FET proteins but no other transportin cargos in FTLD-FUS and is absent in FUS inclusions in ALS with FUS mutations. Acta Neuropathol 124, 705–716 (2012).

30. Rosen, D. R. et al. Mutations in Cu/Zn superoxide dismutase gene are associated with familial amyotrophic lateral sclerosis. Nature 362, 59–62 (1993).

31. Neumann, M. et al. Ubiquitinated TDP-43 in Frontotemporal Lobar Degeneration and Amyotrophic Lateral Sclerosis. Science 314, 130–133 (2006).

32. Rummens, J. et al. TDP-43 seeding induces cytoplasmic aggregation heterogeneity and nuclear loss of function of TDP-43. Neuron 10, 1597–1613.e8 (2025).

33. Scialò, C. et al. Seeded aggregation of TDP-43 induces its loss of function and reveals early pathological signatures. Neuron 113, 1614–1628.e11 (2025).

34. Ling, J. P., Pletnikova, O., Troncoso, J. C. & Wong, P. C. TDP-43 repression of nonconserved cryptic exons is compromised in ALS-FTD. Science 349, 650–655 (2015).

35. Ibrahim, F., et al. Identification of In Vivo, Conserved, TAF15 RNA Binding Sites Reveals the Impact of TAF15 on the Neuronal Transcriptome. Cell Reports 3, 301–308 (2013).

36. Morohoshi, F., Arai, K., Takahashi, E., Tanigami, A. & Ohki, M. Cloning and Mapping of a Human*RBP56*Gene Encoding a Putative RNA Binding Protein Similar to FUS/TLS and EWS Proteins. Genomics 38, 51–57 (1996).

37. Morohoshi, F. et al. Genomic structure of the human *RBP56/hTAFII68* and *FUS/TLS* genes. Gene 221, 191–198 (1998).

38. Bertolotti, A., Lutz, Y., Heard, D. J., Chambon, P. & Tora, L. hTAF(II)68, a novel RNA/ssDNA-binding protein with homology to the pro-oncoproteins TLS/FUS and EWS is associated with both TFIID and RNA polymerase II. The EMBO Journal 15, 5022–5031 (1996).

39. Shi, Y. et al. Structure-based classification of tauopathies. Nature 598, 359–363 (2021).

40. Konstantoulea, K. et al. TAF15 amyloids propagate via defined motifs in a prion-like fashion. 2025.11.17.688886 Preprint at 10.1101/2025.11.17.688886 (2025).

41. Shi, Y. et al. Cryo-EM structures of tau filaments from Alzheimer’s disease with PET ligand APN-1607. Acta Neuropathol 141, 697–708 (2021).

42. Merz, G. E. et al. Stacked binding of a PET ligand to Alzheimer’s tau paired helical filaments. Nat Commun 14, 3048 (2023).

43. Shi, Y., Ghetti, B., Goedert, M. & Scheres, S. H. W. Cryo-EM Structures of Chronic Traumatic Encephalopathy Tau Filaments with PET Ligand Flortaucipir. J Mol Biol 435, 168025 (2023).

44. Kunach, P. et al. Cryo-EM structure of Alzheimer’s disease tau filaments with PET ligand MK-6240. Nat Commun 15, 8497 (2024).

45. Murray, K. A. et al. De novo designed protein inhibitors of amyloid aggregation and seeding. Proc Natl Acad Sci USA 119, e2206240119 (2022).

46. Seidler, P. M. et al. Structure-based discovery of small molecules that disaggregate Alzheimer’s disease tissue derived tau fibrils in vitro. Nat Commun 13, 5451 (2022).

47. Ticozzi, N. et al. Mutational analysis reveals the FUS homolog TAF15 as a candidate gene for familial amyotrophic lateral sclerosis. Am J Med Genet B Neuropsychiatr Genet 156, 285–290 (2011).

48. Couthouis, J. et al. A yeast functional screen predicts new candidate ALS disease genes. Proc Natl Acad Sci USA 108, 20881–20890 (2011).

49. Van Daele, S. H. et al. Genetic variability in sporadic amyotrophic lateral sclerosis. Brain 146, 3760–3769 (2023).

50. Neumann, M. & Mackenzie, I. R. FUS/FET proteinopathies. in Greenfield’s Neuropathology 10e Set (CRC Press, 2024).

51. Li, H. Aligning sequence reads, clone sequences and assembly contigs with BWA-MEM. Preprint at 10.48550/arXiv.1303.3997 (2013).

52. Poplin, R. et al. Scaling accurate genetic variant discovery to tens of thousands of samples. 201178 Preprint at 10.1101/201178 (2018).

53. Danecek, P. et al. Twelve years of SAMtools and BCFtools. Gigascience 10, giab008 (2021).

54. McLaren, W. et al. The Ensembl Variant Effect Predictor. Genome Biol 17, 122 (2016).

55. Zivanov, J., Nakane, T. & Scheres, S. H. W. A Bayesian approach to beam-induced motion correction in cryo-EM single-particle analysis. IUCrJ 6, 5–17 (2019).

56. Rohou, A. & Grigorieff, N. CTFFIND4: Fast and accurate defocus estimation from electron micrographs. J Struct Biol 192, 216–221 (2015).

57. Kimanius, D., Dong, L., Sharov, G., Nakane, T. & Scheres, S. H. W. New tools for automated cryo-EM single-particle analysis in RELION-4.0. Biochem J 478, 4169–4185 (2021).

58. Scheres, S. H. W. Amyloid structure determination in RELION –3.1. Acta Crystallogr D Struct Biol 76, 94–101 (2020).

59. Zivanov, J., Nakane, T. & Scheres, S. H. W. Estimation of high-order aberrations and anisotropic magnification from cryo-EM data sets in RELION-3.1. IUCrJ 7, 253–267 (2020).

60. Chen, S. et al. High-resolution noise substitution to measure overfitting and validate resolution in 3D structure determination by single particle electron cryomicroscopy. Ultramicroscopy 135, 24–35 (2013).

61. Casañal, A., Lohkamp, B. & Emsley, P. Current developments in Coot for macromolecular model building of Electron Cryo-microscopy and Crystallographic Data. Protein Sci 29, 1055–1064 (2020).

62. Croll, T. I. ISOLDE: a physically realistic environment for model building into low-resolution electron-density maps. Acta Cryst D 74, 519–530 (2018).

63. Brown, A. et al. Tools for macromolecular model building and refinement into electron cryo-microscopy reconstructions. Acta Cryst D 71, 136–153 (2015).

64. Yamashita, K., Palmer, C. M., Burnley, T. & Murshudov, G. N. Cryo-EM single-particle structure refinement and map calculation using Servalcat. Acta Cryst D 77, 1282–1291 (2021).

65. Liebschner, D. et al. Macromolecular structure determination using X-rays, neutrons and electrons: recent developments in Phenix. Acta Cryst D 75, 861–877 (2019).

66. Williams, C. J. et al. MolProbity: More and better reference data for improved all-atom structure validation. Protein Sci 27, 293–315 (2018).

