## Supplementary Data for "Distinct TAF15 amyloid filament folds define multiple subtypes of FTLD-TAF15"

**SUPPLEMENTARY FIGURES AND TABLES**

**Supplementary Figure 1: Neuropathological characterisation of NIFID and BIBD cases. a,** Haematoxylin (blue) and eosin (pink) (H&E) staining (left column) and immunohistochemistry (brown) for FUS (middle column) and $\alpha$-internexin (right column) in prefrontal cortex sections from NIFID cases 4, 6, and 9-11, and BIBD cases 1-3. Sections for immunohistochemistry were counterstained with haematoxylin (blue). NIFID cases were characterised by Pick-body like and hyaline conglomerate NCIs visible by H&E stains, FUS-immunoreactive inclusions, and $\alpha$-internexin-immunoreactive inclusions. BIBD cases were characterised by basophilic inclusions visible by H&E stains and FUS-immunoreactive inclusions, in the absence of $\alpha$-internexin-immunoreactive inclusions. FUS-immunoreactive inclusions in NIFID cases 1-8, which had TAF15 filament fold B, were predominantly compact round NCIs and neuropil dots. FUS-immunoreactive inclusions in NIFID cases 9-11 and BIBD cases 1-3, which had TAF15 filament folds C-H, respectively, were predominantly annular and tangle-like NCIs and dystrophic neurites. Scale bars, 25 μm. **b,** Immunoblots for FUS, EWS and TAF15 of representative sarkosyl-insoluble extracts from the prefrontal cortex of aFTLD-U, NIFID and BIBD cases. Case numbers are indicated.

**Supplementary Figure 2: Overview of cryo-EM structures of TAF15 filaments in FTLD-FET.** Central slices centred on the helical axis of the 32 cryo-EM reconstructions of TAF15 filaments determined in this study from aFTLD-U cases 1-3, NIFID cases 1-11 and BIBD cases 1-3.

**Supplementary Figure 3: Cryo-EM structures of TAF15 filaments with fold A in aFTLD-U cases 1-3. a,** Representative micrograph of TAF15 filaments with fold A. **b,** Reference-free 2D class average encompassing one helical cross-over. **c,** 3D reconstruction viewed along the helical axis, showing well-resolved individual TAF15 molecules. **d,** Local resolution estimate for the cryo-EM reconstruction. **e,** Fourier shell correlation (FSC) curves for the two independently-refined cryo-EM half-maps (black lines); for the refined atomic model against the cryo-EM density map (cyan); for the atomic model shaken and refined using the first half-map against the first half-map (green); and for the same atomic model against the second half-map (orange). FSC thresholds of 0.143 and 0.5 are shown. **f,** 3D reconstructions of TAF15 filaments from aFTLD-U cases 1-3 (outlines) and the atomic model PDB ID 8ONS, shown for a single TAF15 molecule perpendicular to the helical axis. The atomic model is colour-coded as a gradient from N- to C-terminal residues. **g,** 3D reconstructions of TAF15 filaments from aFTLD-U cases 1-3, shown as central slices perpendicular to the helical axis and colour-coded to match the outlines of the 3D reconstructions for the respective cases in **(f)**. **a-c,** Scale bars, 10 nm.

**Supplementary Figure 4: Cryo-EM structures of TAF15 filaments with fold B in NIFID cases 1-8. a,** Representative micrograph of TAF15 filaments with fold B. Filament segment coordinates for variants I (cyan), II (magenta) and III (yellow) are shown on right image. **b-d,** Reference-free 2D class averages encompassing one helical cross-over of TAF15 filament fold B variant I **(b)**, variant II **(c)** and variant III **(d)**. **e-g,** 3D reconstructions viewed along the helical axis of TAF15 filament fold B variant I **(e)**, variant II **(f)** and variant III **(g)**, showing well-resolved individual TAF15 molecules. **h-j,** Local resolution estimates for the cryo-EM reconstructions of TAF15 filament fold B variant I **(h)**, variant II **(i)** and variant III **(j)**. **k-m,** Fourier shell correlation (FSC) curves for the two independently-refined cryo-EM half-maps (black lines); for the refined atomic model against the cryo-EM density map (cyan); for the atomic model shaken and refined using the first half-map against the first half-map (green); and for the same atomic model against the second half-map (orange) of TAF15 filament fold B variant I **(k)**, variant II **(l)** and variant III **(m)**. FSC thresholds of 0.143 and 0.5 are shown. **n,** Graph showing the proportion of each variant (I, green; II, blue; III, red) from each individual (1-8). **a-g,** Scale bars, 10 nm.

**Supplementary Figure 5: Cryo-EM structures of TAF15 filaments with fold C in NIFID case 9. a,** Representative micrographs of singlet and doublet TAF15 filaments with fold C. **b-d,** Reference-free 2D class averages encompassing one helical cross-over of TAF15 filament fold C variant I **(b)**, variant I doublet **(c)** and variant II **(d)**. **e-g,** 3D reconstructions viewed along the helical axis of TAF15 filament fold C variant I **(e)**, variant I doublet **(f)** and variant II **(g)**, showing well-resolved individual TAF15 molecules. **h-j,** Local resolution estimates for the cryo-EM reconstructions of TAF15 filament fold C variant I **(h)**, variant I doublet **(i)** and variant II **(j)**. **k-m,** Fourier shell correlation (FSC) curves for the two independently-refined cryo-EM half-maps (black lines); for the refined atomic model against the cryo-EM density map (cyan); for the atomic model shaken and refined using the first half-map against the first half-map (green); and for the same atomic model against the second half-map (orange) of TAF15 filament fold C variant I **(k)**, variant I doublet **(l)** and variant II **(m)**. FSC thresholds of 0.143 and 0.5 are shown. **n,** Cryo-EM reconstruction (black outline) and atomic model of doublet TAF15 filaments with fold C variant I centred on the filament interface. The atomic model is colour-coded as a gradient from N- to C-terminal residues. Hydrogen bonds are shown as dashed lines. **o,** Overlay of the map of TAF15 filament fold C variant I (blue) with the map and model of variant II (yellow). **a-j,** Scale bars, 10 nm.

**Supplementary Figure 6: Cryo-EM structures of TAF15 filaments with folds D and D' in NIFID case 10. a,** Representative micrograph of TAF15 filaments with folds D and D'. **b and c,** Reference-free 2D class averages encompassing one helical cross-over of TAF15 filament folds D **(b)** and D' **(c)**. **d and e,** 3D reconstructions viewed along the helical axis of TAF15 filament folds D **(d)** and D' **(e)**, showing well-resolved individual TAF15 molecules. **f and g,** Local resolution estimates for the cryo-EM reconstructions of TAF15 filament folds D **(f)** and D' **(g)**. **h and i,** Fourier shell correlation (FSC) curves for the two independently-refined cryo-EM half-maps (black lines); for the refined atomic model against the cryo-EM density map (cyan); for the atomic model shaken and refined using the first half-map against the first half-map (green); and for the same atomic model against the second half-map (orange) of TAF15 filament folds D **(h)** and D' **(i)**. FSC thresholds of 0.143 and 0.5 are shown. **j,** Overlay of the maps of TAF15 filament folds D (teal) and D’ (purple). **a-e,** Scale bars, 10 nm.

**Supplementary Figure 7: Cryo-EM structures of TAF15 filaments with the fold E in NIFID case 11. a,** Representative micrograph of TAF15 filaments with fold E. **b,** Reference-free 2D class averages encompassing one helical cross-over. **c,** 3D reconstruction viewed along the helical axis, showing well-resolved individual TAF15 molecules. **d and e,** Local resolution estimates for the cryo-EM reconstructions of TAF15 filament fold E variant I **(d)** and variant II **(e)**. **f and g,** Fourier shell correlation (FSC) curves for the two independently-refined cryo-EM half-maps (black lines); for the refined atomic model against the cryo-EM density map (cyan); for the atomic model shaken and refined using the first half-map against the first half-map (green); and for the same atomic model against the second half-map (orange) of TAF15 filament fold E variant I **(f)** and variant II **(g)**. FSC thresholds of 0.143 and 0.5 are shown. **h,** Overlay of the map of TAF15 filament fold E variant I (yellow) with the map and model of variant II (cyan). **a-c,** Scale bars, 10 nm.

**Supplementary Figure 8: Cryo-EM structures of TAF15 filaments with folds F-H in BIBD cases 1-3. a-c,** Representative micrographs of TAF15 filament folds F **(a)** fold G **(b)** and fold H **(c)**. **d-f,** Reference-free 2D class averages encompassing one helical cross-over of TAF15 filament folds F **(d)** fold G **(e)** and fold H **(f)**. **g-i,** 3D reconstructions viewed along the helical axis of TAF15 filament folds F **(g)** fold G **(h)** and fold H **(i)**, showing well-resolved individual TAF15 molecules. **j-l,** Local resolution estimates for the cryo-EM reconstructions of TAF15 filament folds F **(j)** fold G **(k)** and fold H **(l)**. **m-o,** Fourier shell correlation (FSC) curves for the two independently-refined cryo-EM half-maps (black lines); for the refined atomic model against the cryo-EM density map (cyan); for the atomic model shaken and refined using the first half-map against the first half-map (green); and for the same atomic model against the second half-map (orange) of TAF15 filament folds F **(m)** fold G **(n)** and fold H **(o)**. FSC thresholds of 0.143 and 0.5 are shown. **a-i,** Scale bars, 10 nm.

**Supplementary Figure 9: Recurrent structural motifs in TAF15 filament folds. a-d,** Overlay of atomic models of the indicated TAF15 filament folds focussed on the following recurrent structural motifs: A boot-shaped hairpin formed by Y56-Y78 **(a)**, a hairpin formed by E71-Q86 **(b)**, a hairpin formed by Y78-S95 **(c)** and a hairpin formed by Y53-Y63 **(d)**. The arrows indicate salt bridges between E71 and K74.

**Supplementary Table 1: Demographic, clinical and genetic data**

AAO, age at onset; AD, Alzheimer's disease; aFTLD-U, atypical frontotemporal lobar degeneration with ubiquitin-positive inclusions; ALS, amyotrophic lateral sclerosis; BIBD, basophilic inclusion body disease; CBS, corticobasal syndrome; bvFTD, behavioural variant frontotemporal dementia; CBS, corticobasal syndrome; DD, disease duration; F, female; M, male; MCI, mild cognitive impairment; NIFID, neuronal intermediate filament disease; PPA, primary progressive aphasia; PSP, progressive supranuclear palsy.

**Supplementary Table 2: Neuropathological characterization and semiquantitative assessment of TAF15-immunoreactive pathology**

aFTLD-U, atypical frontotemporal lobar degeneration with ubiquitin-positive inclusions; BI, basophilic inclusions; BIBD, basophilic inclusion body disease; DN, dystrophic neurites; HC, hyaline conglomerate inclusions; NIFID, neuronal intermediate filament disease; NCI, neuronal cytoplasmic inclusions; PBL, pick-body like inclusions. Grading schemes: #: 0, absent; 1, mild/rare; 2, moderate; 3, numerous/frequent. @: 0, absent; 1, <25% of NCIs; 2, between 25% and 75% of NCIs; 3, >75% of NCIs.

**Supplementary Table 3: Cryo-EM data collection, refinement and validation statistics**

Supplementary Figure 1

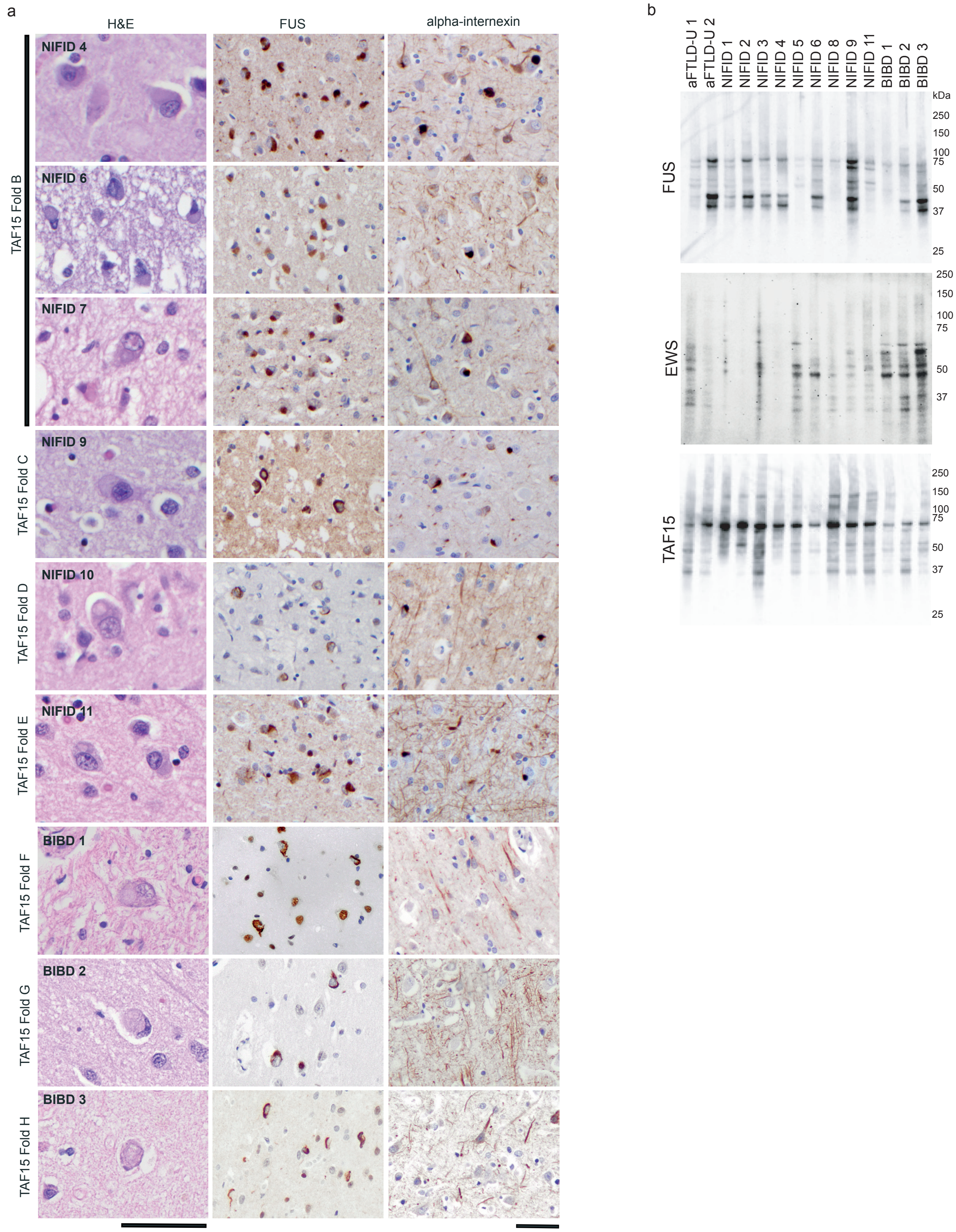

Supplementary Figure 2

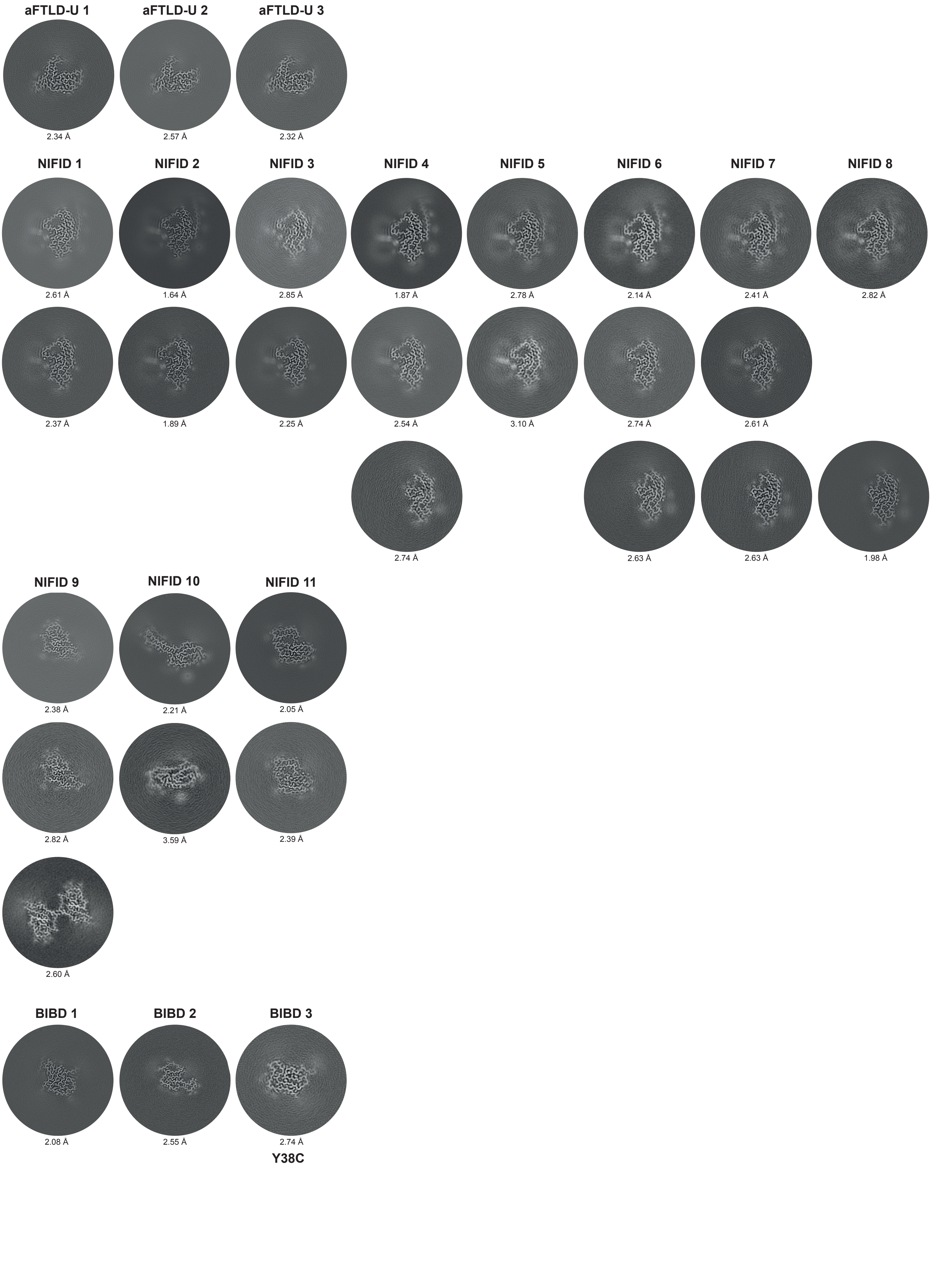

Supplementary Figure 3

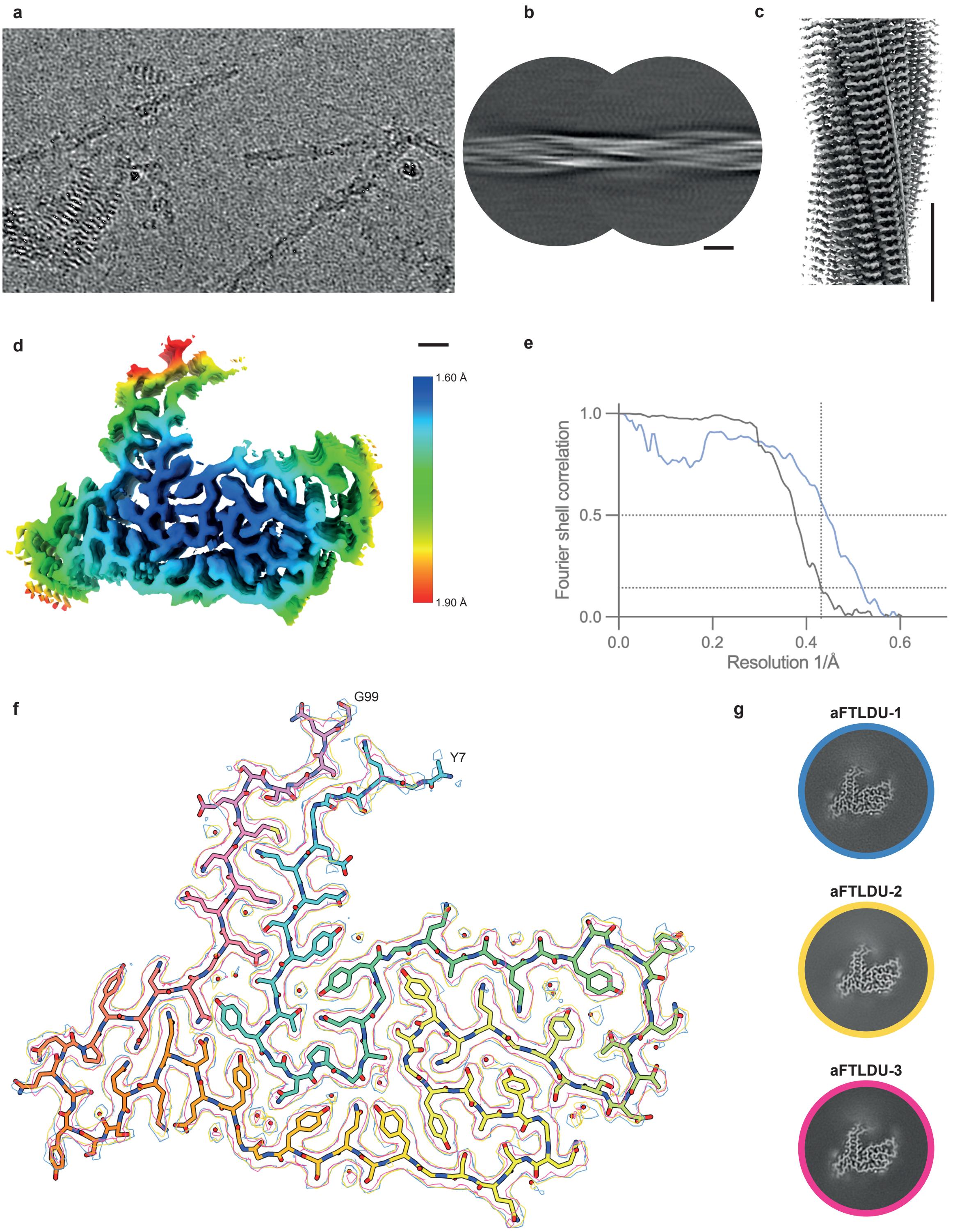

Supplementary Figure 4

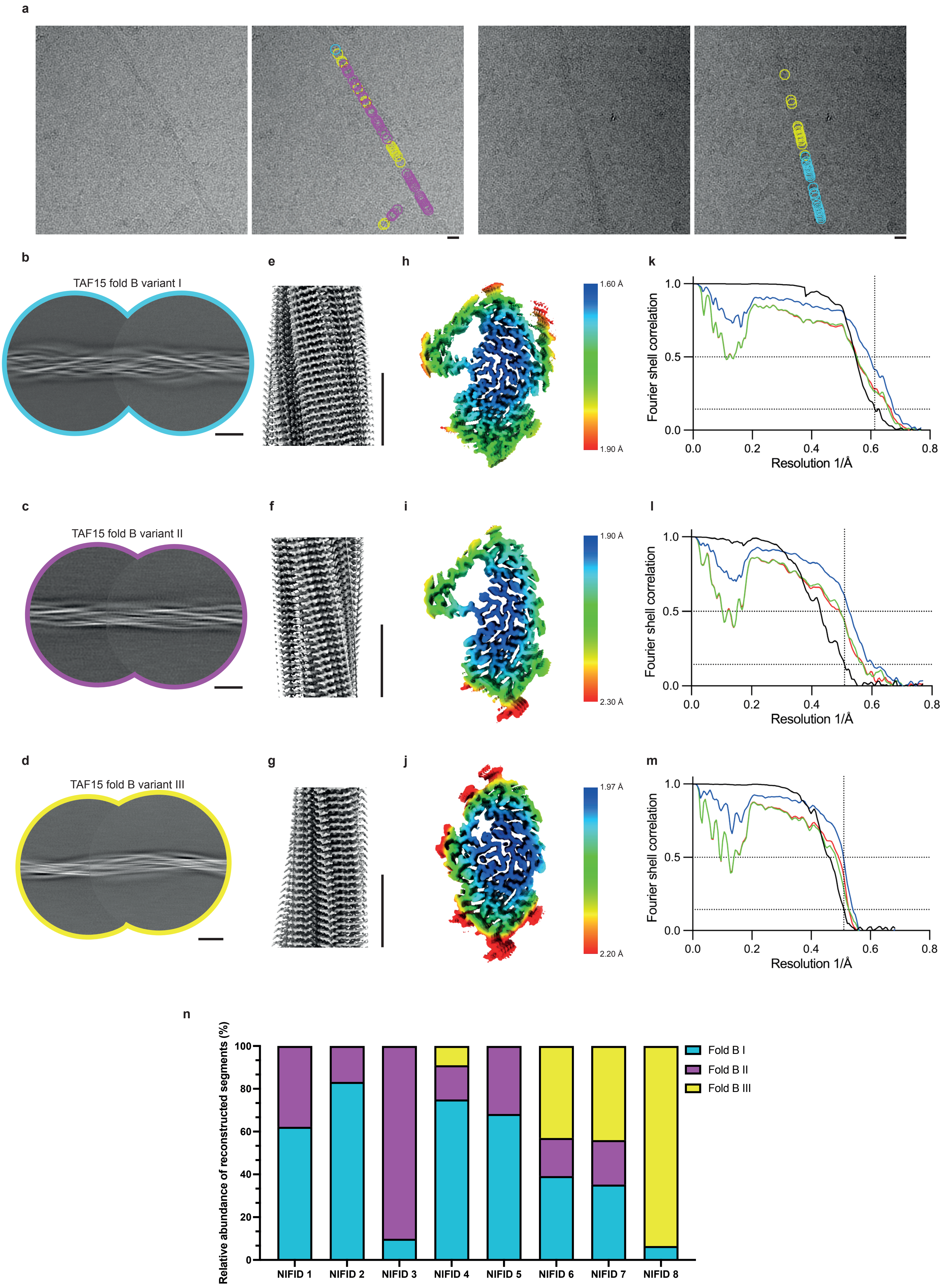

Supplementary Figure 5

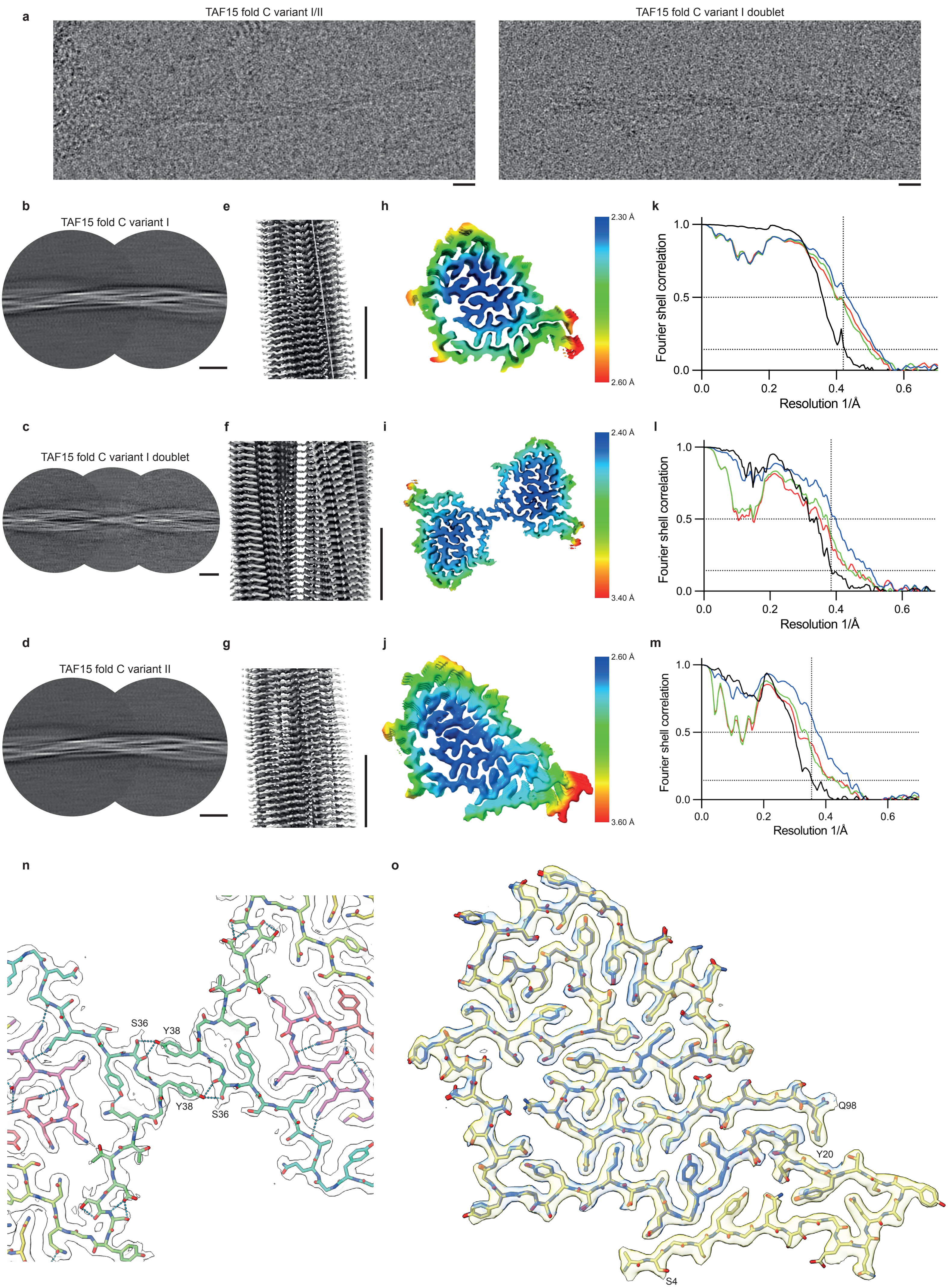

Supplementary Figure 6

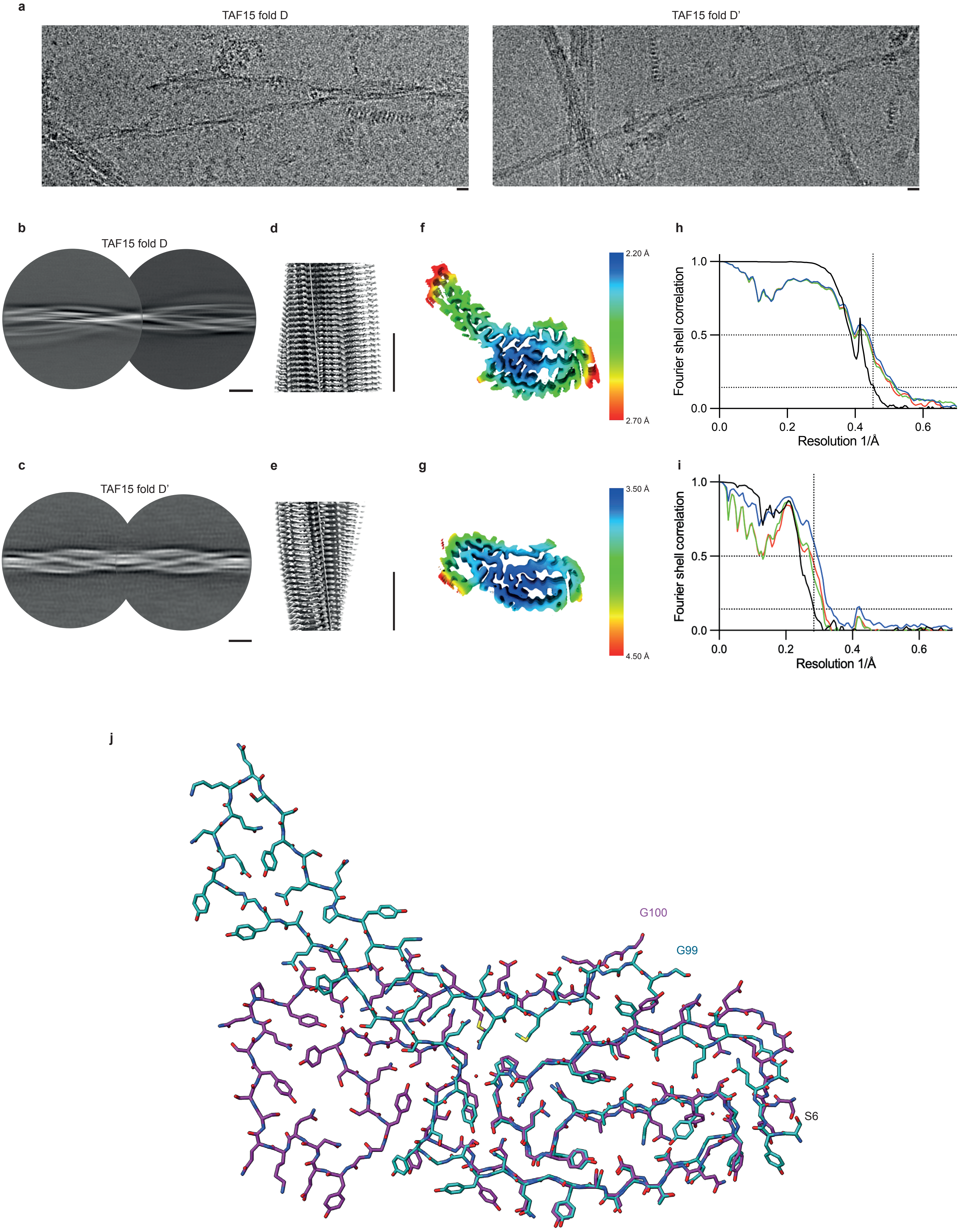

Supplementary Figure 7

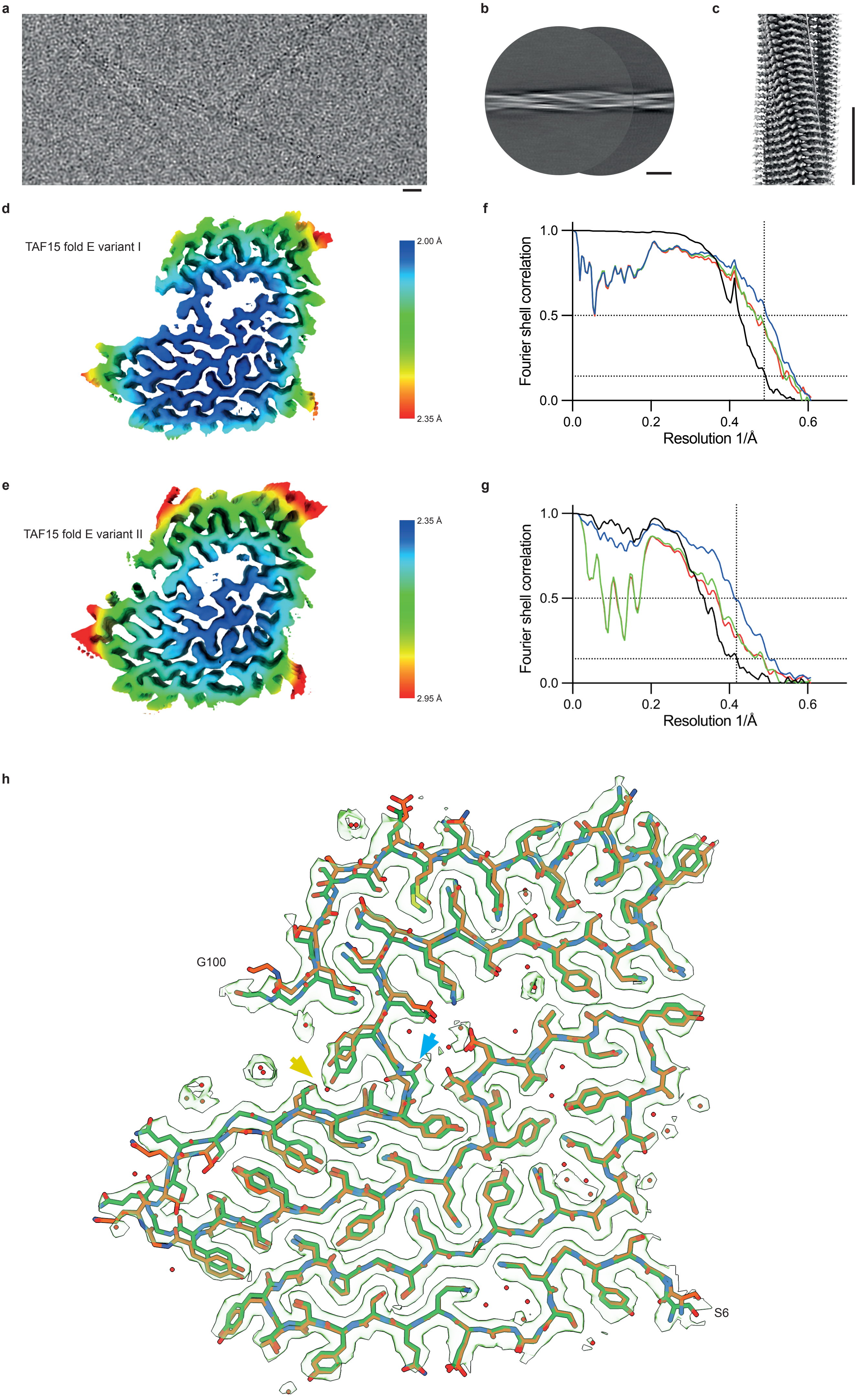

Supplementary Figure 8

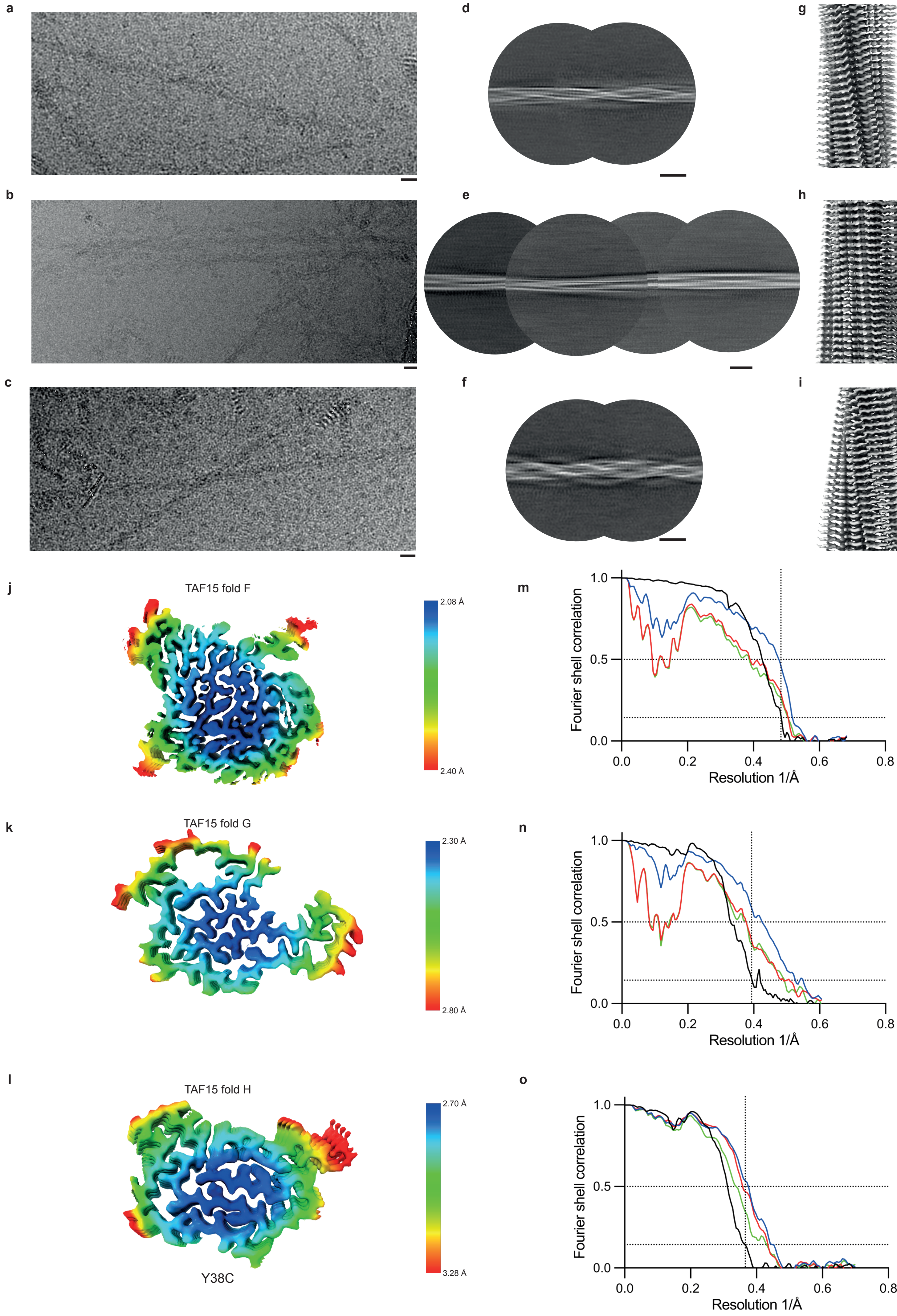

Supplementary Figure 9

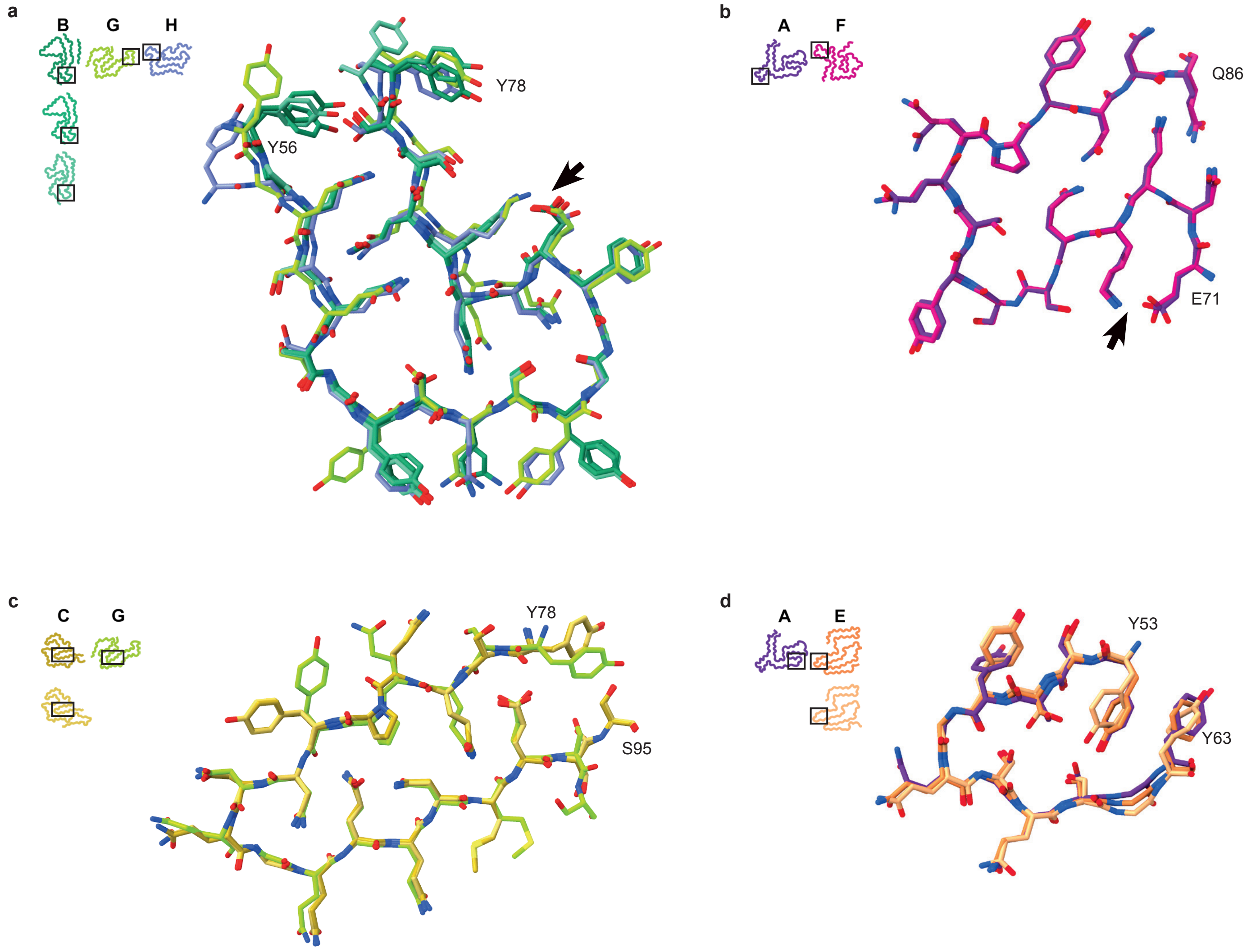

### Supplementary Table 1: Demographic, clinical and genetic data

| Case | M/F | AAO [y] | DD [y] | Clinical diagnosis | *TAF15* |
|---|---|---|---|---|---|
| aFTLD-U 1 | M | 46 | 6 | bvFTD | WT |
| aFTLD-U 2 | F | 36 | 4 | bvFTD | WT |
| aFTLD-U 3 | M | 43 | 10 | bvFTD | WT |
| NIFID 1 | M | 44 | 2 | ALS | WT |
| NIFID 2 | F | 43 | 3 | bvFTD | WT |
| NIFID 3 | F | 41 | 2 | CBS | WT |
| NIFID 4 | F | 36 | 6 | bvFTD, MND | WT |
| NIFID 5 | F | 34 | 7 | PPA, bvFTD | WT |
| NIFID 6 | M | 35 | 3 | bvFTD | WT |
| NIFID 7 | M | 61 | 2 | PPA, PSP, AD | WT |
| NIFID 8 | F | 50 | 4 | CBS | WT |
| NIFID 9 | F | 44 | 2 | bvFTD | WT |
| NIFID 10 | F | 69 | 3 | ALS | WT |
| NIFID 11 | F | 74 | 4 | bvFTD | WT |
| BIBD 1 | F | 43 | 8 | PSP | WT |
| BIBD 2 | F | 70 | 2 | ALS, MCI | WT |
| BIBD 3 | F | 68 | 6 | ALS, PSP | c.113A>G (p.Y38C) |

AAO, age at onset; AD, Alzheimer's disease; aFTLD-U, atypical frontotemporal lobar degeneration with ubiquitin-positive inclusions; ALS, amyotrophic lateral sclerosis; BIBD, basophilic inclusion body disease; CBS, corticobasal syndrome; bvFTD, behavioural variant frontotemporal dementia; CBS, corticobasal syndrome; DD, disease duration; F, female; M, male; MCI, mild cognitive impairment; NIFID, neuronal intermediate filament disease; PPA, primary progressive aphasia; PSP, progressive supranuclear palsy.

### Supplementary Table 2: Neuropathological characterisation and semiquantitative assessment of TAF15 immunoreactive pathology

| Case | Cryo EM | H&E | TAF15 IHC | | | | | α-internexin IHC |
|---|---|---|---|---|---|---|---|---|
| | TAF15 Filament Fold | Inclusions | Overall burden of pathology# | NCI Round/oval @ | NCI annular/ tangles@ | DN# | Dots# | NCI# |
| aFTLD-U 1 | Type A | 0 | 3 | 3 | 1 | 2 | 0 | 0 |
| aFTLD-U 2 | Type A | 0 | 2 | 3 | 1 | 1 | 0 | 0 |
| aFTLD-U 3 | Type A | 0 | 3 | 3 | 1 | 3 | 0 | 0 |
| NIFID 1 | Type B | PBL ++, HC + | 3 | 2 | 2 | 0 | 2 | 3 |
| NIFID 2 | Type B | PBL ++, HC + | 2 | 2 | 2 | 0 | 1 | 2 |
| NIFID 3 | Type B | PBL +++, HC ++ | 3 | 3 | 1 | 0 | 2 | 3 |
| NIFID 4 | Type B | PBL +++, HC ++, NII + | 3 | 3 | 1 | 0 | 3 | 3 |
| NIFID 5 | Type B | PBL +++, HC +, NII + | 3 | 3 | 1 | 1 | 3 | 2 |
| NIFID 6 | Type B | PBL +++, HC + | 3 | 3 | 1 | 0 | 3 | 2 |
| NIFID 7 | Type B | PBL ++, HC + | 3 | 2 | 2 | 0 | 2 | 2 |
| NIFID 8 | Type B | PBL ++, HC +, NII + | 2 | 1 | 3 | 1 | 1 | 3 |
| NIFID 9 | Type C | PBL +, HC +, | 3 | 1 | 3 | 2 | 0 | 1 |
| NIFID 10 | Type D | PBL +, HC + | 2 | 1 | 3 | 2 | 0 | 2 |
| NIFID 11 | Type E | PBL +, HC +, | 3 | 1 | 3 | 3 | 0 | 2 |
| BIBD 1 | Type F | BI + | 1 | 1 | 3 | 1 | 1 | 0 |
| BIBD 2 | Type G | BI + | 2 | 1 | 3 | 1 | 0 | 0 |
| BIBD 3 | Type H | BI ++ | 2 | 0 | 3 | 2 | 0 | 0 |

aFTLD-U, atypical frontotemporal lobar degeneration with ubiquitin-positive inclusions; BI, basophilic inclusions; BIBD, basophilic inclusion body disease; DN, dystrophic neurites; HC, hyaline conglomerate inclusions; NIFID, neuronal intermediate filament disease; NCI, neuronal cytoplasmic inclusions; PBL, pick-body like inclusions
grading schemes:
#: 0, absent; 1, mild/rare; 2, moderate; 3, numerous/frequent
@: 0, absent; 1, <25% of NCIs; 2, between 25% and 75% of NCIs; 3, >75% of NCIs

**Supplementary Table 3: Cryo-EM data collection, refinement and validation statistics**

| TAF15 fold | Type A | | | Type B | | | | | | | | | | | | | | | | | | | | | | | | Type C | | | Type D | | Type E | | Type F | Type G | Type H |
|---|---|---|---|---|---|---|---|---|---|---|---|---|---|---|---|---|---|---|---|---|---|---|---|---|---|---|---|---|---|---|---|---|---|---|---|---|---|
| TAF15 fold variant | - | | | I | | | | | | | | II | | | | | | | | III | | | | | | | | I | II | I, doublet | D | D' | I | II | - | - | - |
| Individual | aFTLD-U 1 | aFTLD-U 2 | aFTLD-U 3 | NIFID 1 | NIFID 2 | NIFID 3 | NIFID 4 | NIFID 5 | NIFID 6 | NIFID 7 | NIFID 8 | NIFID 1 | NIFID 2 | NIFID 3 | NIFID 4 | NIFID 5 | NIFID 6 | NIFID 7 | NIFID 8 | NIFID 1 | NIFID 2 | NIFID 3 | NIFID 4 | NIFID 5 | NIFID 6 | NIFID 7 | NIFID 8 | NIFID 9 | | | NIFID 10 | | NIFID 11 | | BIBD 1 | BIBD 2 | BIBD 3 |
| **Data collection and processing** | | | | | | | | | | | | | | | | | | | | | | | | | | | | | | | | | | | | | |
| Voltage (kV) | 300 | 300 | 300 | 300 | 300 | 300 | 300 | 300 | 300 | 300 | 300 | 300 | 300 | 300 | 300 | 300 | 300 | 300 | - | - | - | - | 300 | - | 300 | 300 | 300 | 300 | | | 300 | | 300 | | 300 | 300 | 300 |
| Electron source | X-FEG | X-FEG | S-FEG | X-FEG | X-FEG | X-FEG | X-FEG | X-FEG | X-FEG | S-FEG | C-FEG | X-FEG | X-FEG | X-FEG | X-FEG | X-FEG | X-FEG | S-FEG | - | - | - | - | X-FEG | - | X-FEG | S-FEG | C-FEG | X-FEG | | | S-FEG | | X-FEG | | X-FEG | C-FEG | S-FEG |
| Detector | Falcon 4i | Falcon 4i | K3 | K3 | Falcon 4i | K3 | Falcon 4i | Falcon 4i | K3 | K3 | Falcon 4i | K3 | Falcon 4i | K3 | Falcon 4i | Falcon 4i | K3 | K3 | - | - | - | - | Falcon 4i | - | K3 | K3 | Falcon 4i | Falcon 4i | | | K3 | | Falcon 4i | | K3 | Falcon 4i | K3 |
| Electron exposure (e⁻/Å $^2$) | 40 | 40 | 40 | 40 | 40 | 40 | 40 | 40 | 40 | 40 | 40 | 40 | 40 | 40 | 40 | 40 | 40 | 40 | - | - | - | - | 40 | - | 40 | 40 | 40 | 40 | | | 40 | | 40 | | 40 | 40 | 40 |
| Defocus range (μm) | -2.0 to -0.5 | -2.5 to -0.5 | -2.0 to -0.5 | -2.0 to -0.5 | -2.0 to -0.5 | -2.0 to -0.5 | -2.0 to -0.5 | -2.0 to -0.5 | -2.0 to -0.5 | -2.0 to -0.5 | -2.0 to -0.5 | -2.0 to -0.5 | -2.0 to -0.5 | -2.0 to -0.5 | -2.0 to -0.5 | -2.0 to -0.5 | -2.0 to -0.5 | -2.0 to -0.5 | - | - | - | - | -2.0 to -0.5 | - | -2.0 to -0.5 | -2.0 to -0.5 | -2.0 to -0.5 | -1.5 to -0.5 | | | -2.5 to -1.0 | | -2.0 to -1.0 | | -2.0 to -1.0 | -1.5 to -0.5 | -2.0 to -1.0 |
| Pixel size (Å) | 0.824 | 0.824 | 0.716 | 0.826 | 0.65 | 0.826 | 0.824 | 0.824 | 0.826 | 0.716 | 0.733 | 0.826 | 0.65 | 0.826 | 0.824 | 0.824 | 0.826 | 0.716 | - | - | - | - | 0.824 | - | 0.826 | 0.716 | 0.733 | 0.65 | | | 0.716 | | 0.824 | | 0.733 | 0.826 | 0.716 |
| Micrographs collected | 14,663 | 31,533 | 17,437 | 36,183 | 15,741 | 12,063 | 13,649 | 14,271 | 34,974 | 15,786 | 16,344 | 36,183 | 15,741 | 12,063 | 13,649 | 14,271 | 34,974 | 15,786 | - | - | - | - | 13,649 | - | 34,974 | 15,786 | 16,344 | 36,121 | | | 56,489 | | 32,977 | | 39,715 | 104,625 | 15,290 |
| Initial particle images (no.) | 135,616 | 428,865 | 140,321 | 517,327 | 850,270 | 373,828 | 777,778 | 129,114 | 556,720 | 260,918 | 347,868 | 517,327 | 850,270 | 373,828 | 777,778 | 129,114 | 556,720 | 260,918 | - | - | - | - | 777,778 | - | 556,720 | 260,918 | 347,868 | 136,044 | | | 1,111,556 | | 478,902 | | 159,941 | 1,097,717 | 85,300 |
| Final particle images (no.) | 26,054 | 215,704 | 104,810 | 137,381 | 266,924 | 15,861 | 148,824 | 41,916 | 22,582 | 74,198 | 15,106 | 115,862 | 21,609 | 105,234 | 59,898 | 25,231 | 11,191 | 34,662 | - | - | - | - | 13,467 | - | 20,312 | 19,469 | 165,317 | 71,272 | 2,600 | 9,846 | 911,798 | 10,168 | 129,198 | 5,728 | 40,188 | 12,437 | 12,482 |
| Symmetry imposed | C1 | C1 | C1 | C1 | C1 | C1 | C1 | C1 | C1 | C1 | C1 | C1 | C1 | C1 | C1 | C1 | C1 | C1 | - | - | - | - | C1 | - | C1 | C1 | C1 | C1 | C1 | C2 | C1 | C1 | C1 | C1 | C1 | C1 | C1 |
| Helical twist (°) | -2.16 | -2.16 | -2.14 | -1.92 | -1.95 | -1.93 | -1.95 | -1.90 | -1.95 | -1.95 | -1.95 | -1.44 | -1.44 | -1.43 | -1.46 | -1.83 | -1.45 | -1.48 | - | - | - | - | -1.53 | - | -1.51 | -1.53 | -1.51 | -1.52 | -1.48 | -1.44 | -1.15 | -2.07 | -1.84 | -1.82 | -1.45 | -0.66 | -2.41 |
| Helical rise (Å) | 4.81 | 4.81 | 4.79 | 4.79 | 4.81 | 4.80 | 4.82 | 4.84 | 4.77 | 4.84 | 4.80 | 4.81 | 4.80 | 4.80 | 4.84 | 4.86 | 4.79 | 4.81 | - | - | - | - | 4.81 | - | 4.78 | 4.84 | 4.79 | 4.83 | 4.83 | 4.79 | 4.77 | 4.81 | 4.83 | 4.84 | 4.80 | 4.83 | 4.79 |
| Map resolution (Å) | 2.34 | 2.57 | 2.32 | 2.61 | 1.64 | 2.85 | 1.87 | 2.78 | 2.14 | 2.41 | 2.82 | 2.37 | 1.96 | 2.25 | 2.54 | 3.10 | 2.74 | 2.61 | - | - | - | - | 2.74 | - | 2.63 | 2.26 | 1.98 | 2.38 | 2.56 | 2.60 | 2.21 | 3.59 | 2.05 | 2.39 | 2.08 | 2.55 | 2.74 |
| FSC threshold | 0.143 | 0.143 | 0.143 | 0.143 | 0.143 | 0.143 | 0.143 | 0.143 | 0.143 | 0.143 | 0.143 | 0.143 | 0.143 | 0.143 | 0.143 | 0.143 | 0.143 | 0.143 | - | - | - | - | 0.143 | - | 0.143 | 0.143 | 0.143 | 0.143 | 0.143 | 0.143 | 0.143 | 0.143 | 0.143 | 0.143 | 0.143 | 0.143 | 0.143 |
| Map resolution range (Å) | 2.22 to 7.36 | 2.58 to 4.85 | 2.27 to 4.58 | 2.52 to 4.34 | 1.62 to 2.06 | 2.78 to 5.61 | 1.82 to 3.19 | 2.54 to 5.60 | 2.10 to 4.31 | 2.30 to 4.74 | 2.43 to 5.59 | 2.33 to 4.31 | 1.90 to 2.72 | 2.12 to 3.66 | 2.41 to 3.58 | 3.15 to 7.73 | 2.64 to 3.94 | 2.48 to 5.23 | - | - | - | - | 2.67 to 4.42 | - | 2.34 to 3.55 | 2.23 to 4.41 | 1.94 to 2.76 | 2.28 to 3.47 | 2.63 to 4.79 | 2.42 to 4.19 | 2.13 to 3.62 | 3.45 to 6.22 | 2.09 to 5.12 | 2.38 to 6.24 | 2.08 to 5.46 | 2.31 to 4.96 | 2.80 to 7.51 |
| **Refinement** | | | | | | | | | | | | | | | | | | | | | | | | | | | | | | | | | | | | | |
| Model resolution (Å) | - | - | - | - | 1.64 | - | - | - | - | - | - | - | 1.96 | - | - | - | - | - | - | - | - | - | - | - | - | - | 1.98 | 2.38 | 2.56 | 2.60 | 2.21 | 3.59 | 2.05 | 2.39 | 2.08 | 2.55 | 2.74 |
| FSC threshold | - | - | - | - | 0.5 | - | - | - | - | - | - | - | 0.5 | - | - | - | - | - | - | - | - | - | - | - | - | - | 0.5 | 0.5 | 0.5 | 0.5 | 0.5 | 0.5 | 0.5 | 0.5 | 0.5 | 0.5 | 0.5 |
| Map sharpening B factor (Å $^2$) | - | - | - | - | -25.9 | - | - | - | - | - | - | - | -22.4 | - | - | - | - | - | - | - | - | - | - | - | - | - | -43.55 | -51.58 | -24.38 | -31.49 | -68.21 | -64.24 | -40.99 | -25.55 | -33.26 | -31.27 | -45.68 |
| Model composition | | | | | | | | | | | | | | | | | | | | | | | | | | | | | | | | | | | | | |
| Non-hydrogen atoms | - | - | - | - | 769 | - | - | - | - | - | - | - | 698 | - | - | - | - | - | - | - | - | - | - | - | - | - | 694 | 603 | 721 | 603 | 715 | 714 | 739 | 746 | 633 | 661 | 642 |
| Protein residues | - | - | - | - | 97 | - | - | - | - | - | - | - | 88 | - | - | - | - | - | - | - | - | - | - | - | - | - | 87 | 79 | 95 | 79 | 94 | 94 | 97 | 97 | 82 | 87 | 85 |
| Solvent atoms | - | - | - | - | 22 | - | - | - | - | - | - | - | 19 | - | - | - | - | - | - | - | - | - | - | - | - | - | 19 | 0 | 0 | 0 | 0 | 0 | 15 | 22 | 0 | 0 | 0 |
| B factors (Å $^2$) | | | | | | | | | | | | | | | | | | | | | | | | | | | | | | | | | | | | | |
| Protein residues | - | - | - | - | 32.72 | - | - | - | - | - | - | - | 28.79 | - | - | - | - | - | - | - | - | - | - | - | - | - | 39.16 | 44.56 | 32.38 | 48.17 | 65.59 | 78.937 | 39.5 | 38.13 | 36.92 | 41.04 | 53.06 |
| Solvent atoms | - | - | - | - | 37.08 | - | - | - | - | - | - | - | 36.01 | - | - | - | - | - | - | - | - | - | - | - | - | - | 42.51 | - | - | - | - | - | 49.54 | 48.14 | - | - | - |
| R.m.s. deviations | | | | | | | | | | | | | | | | | | | | | | | | | | | | | | | | | | | | | |
| Bond lengths (Å) | - | - | - | - | 0.003 | - | - | - | - | - | - | - | 0.002 | - | - | - | - | - | - | - | - | - | - | - | - | - | 0.003 | 0.007 | 0.007 | 0.006 | 0.003 | 0.002 | 0.003 | 0.005 | 0.004 | 0.003 | 0.006 |
| Bond angles (°) | - | - | - | - | 1.017 | - | - | - | - | - | - | - | 0.956 | - | - | - | - | - | - | - | - | - | - | - | - | - | 1.15 | 1.513 | 1.566 | 1.437 | 1.046 | 0.488 | 1.106 | 1.038 | 1.481 | 0.5583 | 0.5383 |
| Validation | | | | | | | | | | | | | | | | | | | | | | | | | | | | | | | | | | | | | |
| MolProbity score | - | - | - | - | 0.61 | - | - | - | - | - | - | - | 0.71 | - | - | - | - | - | - | - | - | - | - | - | - | - | 1.13 | 0.94 | 1.23 | 1.34 | 0.87 | 1.41 | 0.7 | 0.98 | 1.45 | 1.06 | 1.15 |
| Clashscore | - | - | - | - | 0.30 | - | - | - | - | - | - | - | 0.65 | - | - | - | - | - | - | - | - | - | - | - | - | - | 3.39 | 1.82 | 4.2 | 4.44 | 1.16 | 6.73 | 0.51 | 0.57 | 3.12 | 2.7 | 2.73 |
| Favoured rotamers (%) | - | - | - | - | 97.44 | - | - | - | - | - | - | - | 98.59 | - | - | - | - | - | - | - | - | - | - | - | - | - | 98.59 | 98.44 | 97.37 | 96.88 | 100 | 98.67 | 98.68 | 98.68 | 99.71 | 98.57 | 98.53 |
| Poor rotamers (%) | - | - | - | - | 0 | - | - | - | - | - | - | - | 0 | - | - | - | - | - | - | - | - | - | - | - | - | - | 0 | 0 | 0 | 0 | 0 | 0 | 0 | 0 | 0 | 0 | 0 |
| Ramachandran plot | | | | | | | | | | | | | | | | | | | | | | | | | | | | | | | | | | | | | |
| Favoured (%) | - | - | - | - | 98.92 | - | - | - | - | - | - | - | 98.85 | - | - | - | - | - | - | - | - | - | - | - | - | - | 98.84 | 98.7 | 97.87 | 97.4 | 97.83 | 97.85 | 97.89 | 95.79 | 95.00 | 98.82 | 97.59 |
| Allowed (%) | - | - | - | - | 100 | - | - | - | - | - | - | - | 100 | - | - | - | - | - | - | - | - | - | - | - | - | - | 100 | 100 | 100 | 100 | 100 | 100 | 100 | 100 | 100 | 100 | 100 |
| Outliers (%) | - | - | - | - | 0 | - | - | - | - | - | - | - | 0 | - | - | - | - | - | - | - | - | - | - | - | - | - | 0 | 0 | 0 | 0 | 0 | 0 | 0 | 0 | 0 | 0 | 0 |
